# Comparing Brain Asymmetries Independently of Brain Size

**DOI:** 10.1101/2021.12.09.471897

**Authors:** Camille Michèle Williams, Hugo Peyre, Roberto Toro, Franck Ramus

## Abstract

Studies examining cerebral asymmetries typically divide the L-R Measure (e.g., Left– Right Volume) by the L+R Measure to obtain an Asymmetry Index (AI). However, contrary to widespread belief, such a division fails to render the AI independent from the L+R Measure and/or from total brain size. As a result, variations in brain size may bias correlation estimates with the AI or group differences in AI. We investigated how to analyze brain asymmetries in to distinguish global from regional effects, and report unbiased group differences in cerebral asymmetries.

We analyzed the extent to which the L+R Measure, Total Cerebral Measure (TCM, e.g., Total Brain Volume), and L-R TCM predict regional asymmetries. As a case study, we assessed the consequences of omitting each of these predictors on the magnitude and significance of sex differences in asymmetries.

We found that the L+R Measure, the TCM, and the L-R TCM predicted the AI of more than 89% of regions and that their relationships were generally linear. Removing any of these predictors changed the significance of sex differences in 33% of regions and the magnitude of sex differences across 13-42% of regions. Although we generally report similar sex and age effects on cerebral asymmetries to those of previous large-scale studies, properly adjusting for regional and global brain size revealed additional sex and age effects on brain asymmetry.

**Highlights:**

- The typical Asymmetry Index (AI) scales with the size of a region and brain size.
- Omitting the Left+Right Measure influences reported sex differences in asymmetries.
- Omitting brain size or asymmetry influences reported sex differences in asymmetries.
- We report sex and age effects on AIs independent of regional and global brain size.

## 1. Introduction

Symmetry is a central feature of human brain structure. But symmetry is not perfect. Despite global symmetry, numerous neuroanatomical asymmetries have been documented (de Kovel et al., 2017; Ocklenburg et al., 2017; Toga & Thompson, 2003) and appear to have functional significance (Altarelli et al., 2014; Cherbuin et al., 2010; Mazoyer et al., 2014; Zago et al., 2017). These cerebral asymmetries occur across subcortical (Guadalupe et al., 2017), cortical (Kavaklioglu et al., 2017; A. Kong et al., 2017; X.-Z. Kong et al., 2020), and cerebellar (Kavaklioglu et al., 2017; Kurth et al., 2018) volumes, as well as cortical surface areas and thicknesses (X.-Z. Kong et al., 2018).

Brain asymmetries are typically calculated by dividing the difference between the left and right measures (e.g., volume) of a region by their sum: AI = (Left-Right Measure) / (Left+Right Measure), with the measure corresponding to volume, mean thickness, or surface area (X.-Z. Kong et al., 2020; Kurth et al., 2018). Previous studies assumed that dividing by the L+R Measure of a region ensures that the asymmetry index (AI) is independent of the size of this region or brain size (X.-Z. Kong et al., 2020). This adjustment is similar to the proportion method, a method used to adjust for differences in brain size that divides a regional measure (e.g. hippocampal volume) by a Total Cerebral Measure (TCM; e.g., Total Brain Volume (TBV)). However, instead of removing the effect of brain size on the local measure, studies have shown that the proportion approach often inverts the effect of brain size because it assumes that the intercept of the relationship between a local measure and brain size is null and that the relationship is linear^1^ (Lefebvre et al., 2015; Liu et al., 2014; Sanchez-Roige et al., 2019). Since the intercept is generally not null, dividing by the global measure can lead to reporting inaccurate group differences(Reardon et al., 2016; Sanchis-Segura et al., 2019; Williams et al., 2021a, 2021b). To take this issue into account, recent studies adjust for brain size by using the covariate approach, which includes the global measure as a covariate in the linear regression models. This raises the question of whether bilateral (L+R) measures and/or global brain size should also be considered as covariates when examining group differences in asymmetry.

Furthermore, the covariate approach, like the proportion method, assumes that the relationship between a local and a global measure is linear, while a growing literature suggests that the relationships between global and local cerebral measures are non-linear (Finlay et al., 2001; Fish et al., 2017; Liu et al., 2014; Reardon et al., 2016, 2018; Toro et al., 2009). Thus, ignoring non-linearities in the brain could additionally lead to reporting inaccurate group differences.

Finally, global brain asymmetry may also influence local brain asymmetries. Although global brain asymmetries have been investigated across studies linking neuroanatomy to cognition and mental health disorders, the relationship between the global and local AI remains unclear. As global brain asymmetries may partly explain local asymmetries, omitting the L-R TCM could influence reported group differences in asymmetries.

In the present paper, we aimed to identify the brain measures that should be taken into account by studies examining group differences in cerebral asymmetries. We first examined whether the L+R Measure, TCM, and L-R TCM significantly predicted regional asymmetries. We then investigated whether the relationship between the regional AI and the L+R Measure, global brain size, and the L-R TCM were non-linear. Thirdly, since the two sexes differ in brain size, we used the case of sex differences in brain asymmetries to illustrate the consequences of not taking these adjustments into account. Finally, we reported the effects of sex, age, and their interactions when including all necessary brain covariates to report group differences in asymmetries that do not scale with brain size.

## 2. Methods

### 2.1. Participants

Participants were part of the UK Biobank, an open-access large prospective study with phenotypic, genotypic, and neuroimaging data from 500 000 participants recruited between 2006 and 2011 at 40 to 69 years old in Great Britain (Sudlow et al., 2015). All participants provided informed consent (“Resources tab” at https://biobank.ctsu.ox.ac.uk/crystal/field.cgi?id=200). The UK Biobank received ethical approval from the Research Ethics Committee (reference 11/NW/0382) and the present study was conducted under application 46 007.

Imaging-Derived Phenotypes from Magnetic Resonance Imaging (MRI) data are currently available for about 41 000 UK Biobank participants. Imaging-Derived Phenotypes included in this study correspond to volumes, mean thicknesses, surface areas from the first imaging visit and were generated by an image-processing pipeline developed and run by the UK Biobank Imaging team (Alfaro-Almagro et al., 2018; Miller et al., 2016).

#### 2.1.1. Brain Image Acquisition and Processing

A standard Siemens Skyra 3T running VD13A SP4 with a standard Siemens 32-channel RF receive head coil was used to collect data ([Brain Scan Protocol](https://biobank.ctsu.ox.ac.uk/crystal/refer.cgi?id=2367)). The 3D MPRAGE T1-weighted volumes were analyzed by the UK Biobank Imaging team with [pipeline scripts](https://www.fmrib.ox.ac.uk/ukbiobank/) that primarily call for FSL and Freesurfer tools. Details of the acquisition protocols, image processing pipeline, image data files, and derived measures of brain structure and function are available in the [UK Biobank Imaging Protocols](https://biobank.ctsu.ox.ac.uk/crystal/crystal/docs/brain_mri.pdf).

#### 2.1.2. Total Brain Volume (TBV)

TBV was calculated as the sum of the following ASEG segmentation: Cerebellum GMV (Left: 26557, Right: 26588), Cerebral GMV (Left: 26552, Right: 26583), Cerebellum WMV (Left: 26556, Right: 26587), Cerebral WMV (Left: 26553, Right: 26584), and the total subcortical volume we calculated with the ASEG and Freesurfer subfields. Refer to Supplemental Section 1 for details on the choice of TBV. We excluded individuals with missing data in these regions, yielding 40 055 participants.

#### 2.1.3. Scanner Site

The age and sex of participants differed across the 3 scanner sites located in Cheadle (Site 11025), Reading (Site 11026), and Newcastle (Site 11027; Supplemental Section 2). One individual without scanner site location was removed from the analyses, yielding 40 054 participants.

#### 2.1.4. Sex

Participants who did not self-report as male or female or whose self-reported sex and genetic sex differed were also excluded from the analyses (N = 26). When genetic sex was not available, reported sex was used to define the sex of the participant. Of the 40 028 participants included in the analyses, there were more females (N= 21 142) than males (N = 18 886, χ^2^(1) = 127.15, p < 2.2e-16).

#### 2.1.5. Age

To obtain a continuous measure of age, age was calculated based on the year and month of birth of the participant and the day, month, and year of their MRI visit. The mean age was 64.13 years old (*SD* = 7.54). Males (*M* = 64.81 years, *SD* = 7.64) were older than females (*M* = 63.51 years, *SD* =7.39, t _(39186)_ = 17.17, p < 2.2e-16).

#### 2.1.6. Handedness

The age and sex of participants differed between Right-handed (N = 35,596), Left-handed (N = 3,711), and ambidextrous (N = 547) individuals (Supplemental Section 3). Participants without handedness data (N = 174) were excluded from the analyses in Q4.

### 2.2. Brain Measures

The descriptive statistics of all global and regional brain measures analyzed in the present study and their respective data fields and segmentation origin are listed in Supplemental Tables A1-6. Brain measures mainly correspond to GMV since WMVs were not segmented by the UK Biobank Imaging Team.

#### 2.2.1. Global Brain Measures

A total of 10 Global AIs were investigated: Hemispheric Brain Volume, Mean Cortical Thickness, Surface Area, Subcortical GMV, Cortical GMV, Cerebral GMV, Cerebral WMV, 2 measures of Cerebellar GMV (ASEG and FAST FSL), and Cerebellar WMV. We examined two measures of Cerebellar GMV to evaluate the consistency of the FAST FSL and ASEG segmentation algorithms since we previously found sex effects in the opposite direction between the ASEG and the FAST FSL Cerebellar GMVs (Williams et al., 2021).

WMV measures were taken from the left and right Freesurfer ASEG segmentation (data-field 190). The left and right Cerebellum GMV measures correspond to the sum of the left and right cerebellar volumes from the FAST segmentation (data-field 1101) and the left and right ASEG Cerebellum GMVs (Left: data-field 26557, Right: data-field 26588).

Following the recommendations from the [UK Biobank Imaging Protocols](https://biobank.ctsu.ox.ac.uk/crystal/crystal/docs/brain_mri.pdf), we excluded Freesurfer brain measures when T2-FLAIR was not used with the T1 images for the Freesurfer a2009s (volume, surface area, and mean thickness) and Freesurfer subfield segmentations. The left and right Total Mean Cortical Thicknesses (MCT) and Total Surface Area (TSA) respectively corresponded to the left and right mean cortical thickness and surface area measures from the Freesurfer a2009s segmentation (data-field 197). Participants that were missing more than 10% of the mean cortical thicknesses were excluded from the mean thickness analyses. The same criterion was applied to surface areas. The left and right Cortical GMVs correspond to the sum of the left and right volumetric measures from the Freesurfer a2009s segmentation (data-field 197) and the Cerebral GMV corresponds to the ASEG Cerebral GMVs (Left: data-field 26552, Right: data-field 26583). The Subcortical GMV was calculated from the sum of the left and right whole amygdala, hippocampus, and thalamus volumes from the Freesurfer subfields (data-field 191) and the left and right caudate, accumbens, pallidum, ventral diencephalon, and putamen of the Freesurfer ASEG segmentation (data-field 190).

While 40 028 individuals were included in the analyses, 39 238 individuals were included in the FAST segmentation analyses and 38 710 in the Freesurfer a2009s segmentation and subcortical subfields analyses. Missing values and null segmentations (e.g., 0 mm3) for a region were replaced by the mean of that region when calculating global measures.

#### 2.2.2. Regional Brain Measures

A case-wise participant exclusion strategy was applied to each brain measure for the regional analyses: participants with a missing value or a segmentation error for a region were excluded from the analyses of that region but were maintained in the analyses of other brain measures. Following visual examination of the distribution of regional cerebral measures, values 3 times the interquartile range for a region were considered to be segmentation errors and were removed from the analyses of that region as done by Williams and colleagues (2021).

A total of 296 regional AIs were investigated: 222 cortical regions (74 volumes, 74 surface areas, and 74 cortical mean thicknesses) from the Freesurfer a2009s segmentation (Destrieux Atlas, data-field 197), 56 whole segmentation and subfields of the amygdala, hippocampus, and thalamus (Freesurfer subfields, data-field 191), 10 cerebellum grey matter segmentation from the FAST segmentation (data-field 1101), and 8 subcortical and ventricle volumes from the Freesurfer ASEG segmentation (data-field 190). When choices between different types of segmentation were available, we took the smallest segmentation available (e.g. FreeSurfer Tools subfields) to obtain a finer grain analysis.

### 2.3. Statistical Analyses

Analyses were preregistered on OSF and run using R (R Core Team, 2019). The preregistration and code are on OSF (https://osf.io/wt6uf). Packages are listed in Supplemental Section 6. The categorical sex variable was coded -0.5 for males and 0.5 for females. The scanner site was dummy coded with the largest site (Cheadle: Site 1102) as reference and handedness was dummy coded with the right-handed individuals as the reference.

#### Q1: Do the L+R Measure, global brain size, and the L-R TCM predict AI_regional_?

The regional and the global asymmetry index were calculated as AI = (L-R Measure) / (L+R Measure)). We tested whether the L+R Measure, the TCM (TCM), and the L-R TCM significantly predicted the AI by applying equation 1 to each region/global measure. For global measures, the TCM corresponded to Total Brain Volume. For regional measures, the TCM was TBV when examining AIs of volumes, TSA for surface areas, and Total MCT for mean thicknesses.

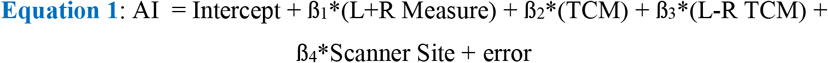

A significant L+R Measure suggested that the regional AI did not properly adjust for the L+R Measure. If any predictor significantly predicted the AI (alpha = 0.05) in more than 5% of regional and global measures, we concluded that the predictor should be included when examining group differences in asymmetry.

Even if dividing by L+R did not properly adjust for L+R, we kept defining the AI as (L-R Measure)/(L+R Measure) instead of defining it as the L-R Measure to obtain a comparable AI across regions.

#### Q2: Is the relationship of the AI with the L+R Measure, TCM, or the L-R TCM, non-linear?

Predictors included in the following models were based on the results from Q1. Here, we provide equations for the case where the L+R Measure, TCM, and the L-R TCM significantly predict 95% or more AIs.

We compared a linear model (equation 1) to standard nonlinear models (equations 2-7): polynomial linear regressions and splines. The comparison was based on the percent change in Residual Standard Error (RSE) between the models^2^. The RSE is the square root of the residual sum of squares divided by the degrees of freedom and is provided in the output of each model using the summary function.

Since more complex models always yield lower RSE, we consider a “significantly better fit” as a reduction of RSE greater than 1%. If the reduction in RSE was larger than 1% in more than 5% of regional and global measures, this justified the use of the non-linear model over the linear one in the subsequent analyses.

The most common splines were selected based on Perperoglou and colleagues’ (2019) review and previous studies examining nonlinearities in neuroimaging studies (e.g., Fjell et al., 2009; Reardon et al., 2018). We used the gam function from the mgcv package (Wood, 2017) to model splines. The s() function will be applied to each predictor in the model for which we want a smoothing parameter.

##### Polynomial Quadratic Equation

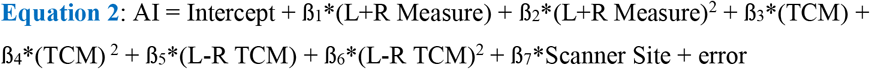

##### Polynomial Cubic Equation

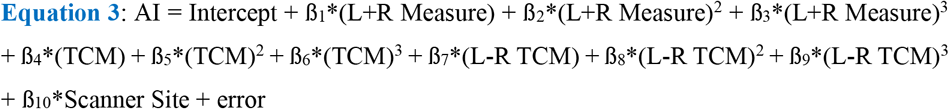

##### B-spline basis Equation, degree of the piecewise polynomial 3 (cubic splines)

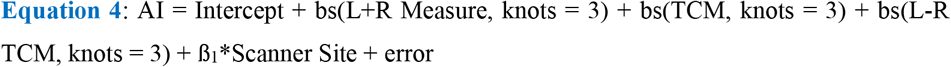

##### Natural spline Equation, degree of the piecewise polynomial 2

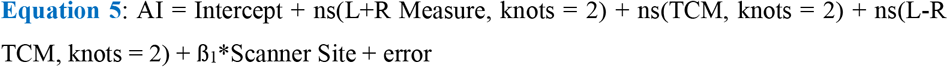

##### Natural spline Equation, degree of the piecewise polynomial 3

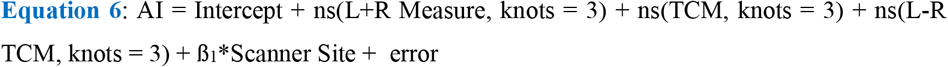

##### Penalized thin plate regression splines

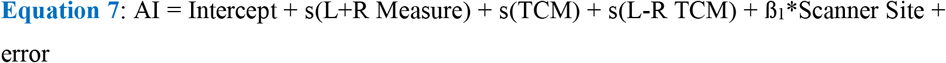

From the results of Q1 and Q2, we determined a “default model for the analysis of brain asymmetries independent of brain size”, including all the predictors determined by Q1, and the best linear or nonlinear model determined by Q2. We then analyzed the consequences of departing from this default model with the case of sex differences in Q3.

#### Q3: To what extent do group estimates change with the included covariates and the type (linear vs. non-linear) of relationship considered between the covariates and AI?

We used sex as our grouping variable and generated equations 8-11 based on the results from Q1 and Q2. For instance, if Q1 showed that the L+R Measure, the TCM, and the L-R TCM were significant predictors of the AIs and Q2 that the reduction in RSE across models was smaller than 1% for 95% or more of the AIs, we ran equation 8 as the default equation and compared it to equations removing one term at a time (equation 9-11). We mean-centered and divided continuous variables by 1 SD to obtain standardized beta estimates.

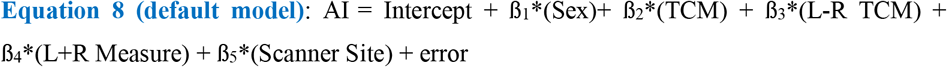

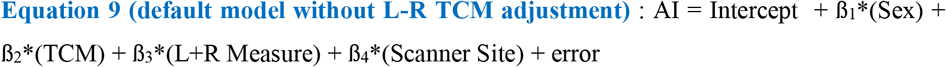

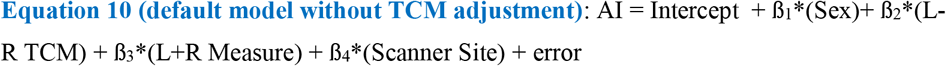

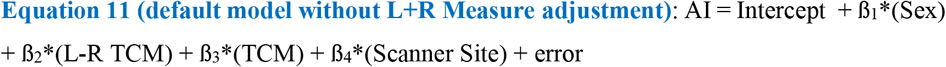

We systematically compared the full model (equation 8) with each subsequent model omitting one type of adjustment. For each pair of models considered, we statistically tested the difference in the sex coefficient ß1 at the alpha=0.05 level by conducting a Z-test, in all regions. For instance, for equations 8 and 9, we computed the Z-score as follows:

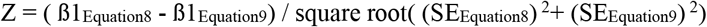

If the |Z-score| was greater or equal to 1.96, then the ß1 of the equations significantly differed at alpha = 0.05. If this test was significant in more than 5% of regional and global measures, we concluded that the omitted covariate led to incorrect estimations of sex differences in asymmetry.

##### Q4: Do cerebral asymmetries vary as a function of age and sex effects and interactions?

We then examined sex differences in asymmetries by applying the model adjustments determined in Q1 and Q2 and by taking into account the effects of age on brain measures, as is commonly done in the literature.

In the case where (i) the L+R Measure, the TCM, and the L-R TCM are significant predictors of AI and (ii) the percent change in RSE is smaller than 1% for 95% or more of the regional and global measures, we examined age (linear and quadratic) and sex effects and age by sex interactions with equation 12. We mean-centered and divided continuous variables by 1 SD to obtain standardized beta estimates. We used polynomials instead of splines to obtain a standardized beta coefficient for non-linear age effects and the non-linear age by sex interactions.

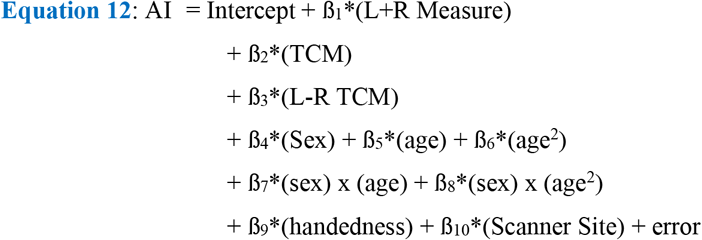

##### Multiple Comparison Corrections

Three thresholds of significance will be provided and will vary depending on the number of cerebral measures (296 regional measures and 10 global measures) and the number of estimates of interests in the performed regressions: α1=0.05/ (5 estimates of interest in regression), α2=0.05/ (306 cerebral measures), and α3=0.05/ (5 estimates of interest in regression * 306 cerebral measures).

##### Additional Analyses

We created in Global Asymmetry Markers, Person-level Asymmetry Deviance Markers, Global Deviance Markers, tested the Sex Difference in the Variance of the regional AIs, and compared our raw AIs and sex and age effects to previous large-scale studies in the Supplemental Section 5.

## 3. Results

### Q1: Do the L+R Measure, TCM size, and the L-R TCM predict the AI?

The L+R Measure and the TCM significantly predicted 89% of global and regional AIs and the L-R TCM significantly predicted 91% of global and regional AIs (respectively, 271/306, 277/305, and 270/305). Estimates and p-values are reported in Supplemental Tables B1-4.

Across global measures, where TBV was the TCM, the L+R TCM predicted the asymmetry in TBV, and the L+R Measure, the L–R TCM, and TCM predicted asymmetries across all other global measures.

The L+R Measure significantly predicted the regional AI in 87% (128/148) of volumes, in 89% (66/74) of surface areas, and in 91% (91/74) of mean thicknesses. The L-R TCM significantly predicted the regional AI in 91% (135/148) of volumes, 81% (60/74) of surface areas, and 99% (73/74) of mean thicknesses. Finally, the TCM significantly predicted the regional AI in 85% (126/148) of volumes, 93% (69/74) of surface areas, and 89% (66/74) of mean thicknesses.

Since the L+R Measure, the L-R TCM, and the TCM were significant predictors of asymmetry in more than 5% of the investigated global and regional AIs, we concluded that the above predictors should be included when examining group differences in asymmetry and did so in the subsequent analyses.

### Q2: Is the relationship of the AI with the L+R Measure, or with global brain size, or with the L-R TCM, non-linear?

The reduction in RSE was only greater than 1% in 2% of regions (6/306). The reduction in RSE was the greatest between the linear model and the smooth thin-plate spline (Supplemental Tables C1-4, Summary in C5). Since the reduction in RSE was more than 1% in less than 5% of the investigated global and regional AIs, we concluded that linear models were sufficient to model the relationship between AI and the L+R Measure, global brain size (e.g. TBV), and the L-R TCM, which were negligibly non-linear. We conducted exploratory analyses to identify the cerebral predictors responsible for the reduction in RSE in the six regions that had a reduction in RSE that was larger than 1% (Supplemental Section 4).

### Q3: To what extent do group estimates change with the included covariates and the type (linear vs. non-linear) of relationship considered between the covariates and AI?

We examined whether group differences change as a function of the predictors included in the model by comparing the magnitude and significance of the standardized sex betas obtained from equations 8 to 11 (Supplemental Tables D1-4). Removing either the L+R Measure or the TCM significantly changed the magnitude of the sex differences in asymmetry (p < 0.05) in 42 % of regions (130/306 and 129/305, respectively). Removing the L-R TCM also significantly changed the magnitude of the sex differences in asymmetry (p < 0.05) but only in 13% of regions (39/305).

When looking at these results for volumes, mean thicknesses, and surface areas, separately, the estimate of the sex differences in asymmetry significantly changed in 49% (72/148) of volumes when omitting TBV, in 7% (11/148) of volumes when omitting the TBV AI, and in 9% (13/148) of volumes when omitting the L+R Measure. The estimate of sex differences in asymmetry significantly changed in 69% (51/74) of surface areas when omitting TSA and in 78% (58/74) of surface areas when omitting the L+R Measure. And the estimate of sex differences in asymmetry significantly changed in 34% (25/74) of mean thicknesses when omitting the Total MCT AI and in 69% (51/74) of surface areas when omitting the L+R Measure. The magnitude of the sex difference in asymmetry did not change when removing the TSA AI from the model on surface areas or when removing Total MCT from the model on mean thicknesses. Removing a cerebral predictor changed the significance of the sex differences in asymmetry (p <0.05) in about 1/3rd of the investigated global and regional measures (Supplemental Tables D5). We additionally report the R2, AIC, and BIC of equations 8-11 and find that the full model is more appropriate across a majority of regions (Supplemental Tables E1-4).

Therefore, we conclude that removing any of the three covariates (L+R Measure, TCM, or L-R TCM) from a model investigating sex differences in regional asymmetries substantially changes the reported findings.

### Q4: Do cerebral asymmetries vary as a function of age and sex effects and interactions?

We preregistered that we would include cerebral predictors in the models examining age and sex effects and interactions based on the results from Q1 and Q2. Therefore, although removing TSA AI for surface areas and Total MCT for mean thicknesses did not change the magnitude of group differences in Q3, these covariates were still included in the Q4 models.

We report age and sex interactions in the main text that survived the strictest Bonferroni correction (p < 0.05 / (5*306)). The effects of all predictors across all regions are available in Supplemental Tables SD1-4. Plots for significant interactions are available in Supplemental File S5. We additionally report the effect of handedness.

Across global and regional measures, there were sex differences in AI in 42% (129/306) of regions, linear age effects in AI in 35% (107/306) of regions, quadratic age effects in AI in 8% (25/306) of regions, linear age by sex interactions in 3 % (10/306) of regions, and no quadratic age by sex interactions.

#### 1. Global Measures

The AI of Cortical GMV and Total MCT linearly increased with age (β = 0.02 and β = 0.03, respectively), whereas the AI of TSA, Total Subcortical Volumes, Cerebral WMV and GMV, Cerebellum GMV (FAST and ASEG) and WMV, and TBV linearly decreased with age. A quadratic age effect was only present for the L-R FAST Cerebellar GMV (β = 0.04). There were no sex differences in the AI of Cerebellar WMV and Total Subcortical Volume. Males had a greater AI than females in terms of Cortical GMV (β = -0.08) and Total MCT (β = -0.11). Finally, females had a greater AI in TSA, Cerebellar GMV (ASEG), Cerebral WMV and GMV, and TBV (Supplemental Table F1).

#### 2. Cortical Regions

The AI was larger in females compared to males in 12% (9/74) of cortical volumes, 19% (14/74) of surface areas, and 22% (16/74) of mean thicknesses. The AI was larger in males compared to females in 24% (18/74) of cortical volumes, 20% (15/74) of surface areas, and 19% (14/74) of mean thicknesses. The sex difference in AI ranged from -0.25 (temporal pole) to 0.14 (superior occipital sulcus and transverse occipital sulcus) for volumes, from -0.20 (posterior ramus of the lateral sulcus) to 0.13 (long insular gyrus and central sulcus of the insula) for surface areas, and from -0.17 (orbital gyrus) to 0.12 (posterior ramus of the lateral sulcus) for mean thicknesses (Figure 1).

**Figure 1.**
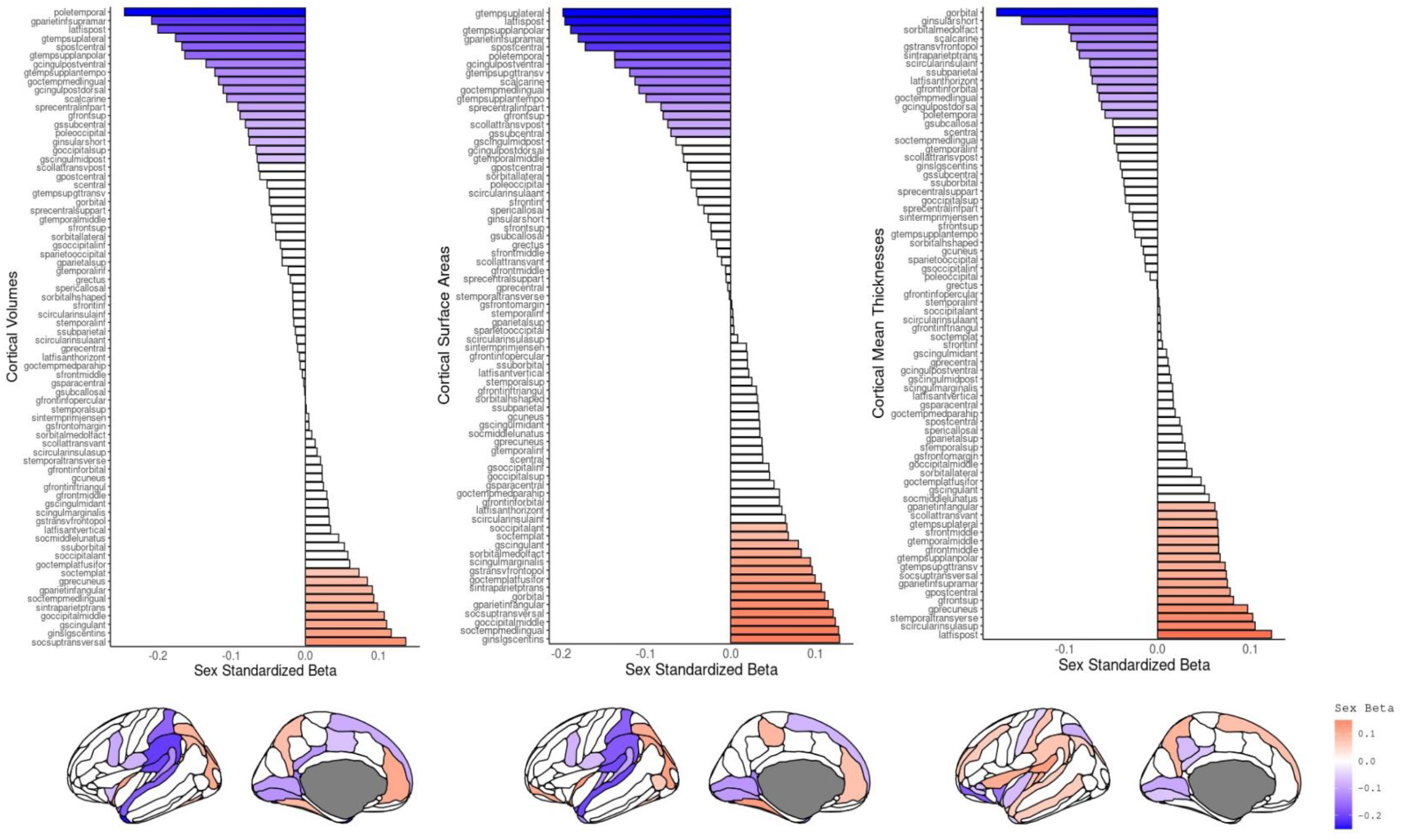
Sex Differences in Asymmetry Index (AI) across Cortical Volumes, Surface Areas, and Mean Thicknesses. Positive sex standardized beta indicates that females have a greater AI than males (red) and a negative slope indicates that males have a greater AI than females (blue). Colors indicate significance at p<0.05/(5*306). Figures made with ggseg (Mowinckel & Vidal-Piñeiro, 2019).

The AIs of males and females were very highly correlated (r>=0.98; Supplemental Information Figure S1-2) and regions with significant sex differences had asymmetries in the same direction across sexes (Supplemental Tables SG1-4) with some exceptions. Males had a leftward asymmetry, whereas females had a rightward asymmetry in the medial orbital sulcus mean thickness (β = -0.10), the posterior-ventral part of the cingulate gyrus surface area (β = - 0.14), and the inferior part of the precentral sulcus (β = -0.09) volume. Females had a leftward asymmetry, whereas males had a rightward asymmetry in the volumes of the long insular gyrus and central sulcus of the insula(β = 0.12).

The AI increased with linear age in 12% (9/74) of cortical volumes, 5% (4/74) of surface areas, and 18% (13/74) of mean thicknesses. The AI decreased with linear age in 10% (7/74) of cortical volumes, 8% (5/74) of surface areas, and 15% (11/74) of mean thicknesses (Figure 2). The AI decreased with quadratic age in the volume of the cuneus gyrus (β = -0.02), in the surface area of the short insular gyri (β = 0.02), and in the mean thicknesses of the suborbital sulcus (β = 0.03) and the inferior temporal gyrus (β = 0.02). There were no significant interactions of age by sex or age^2^ by sex across cortical volumes, mean thicknesses, and surface areas.

**Figure 2.**
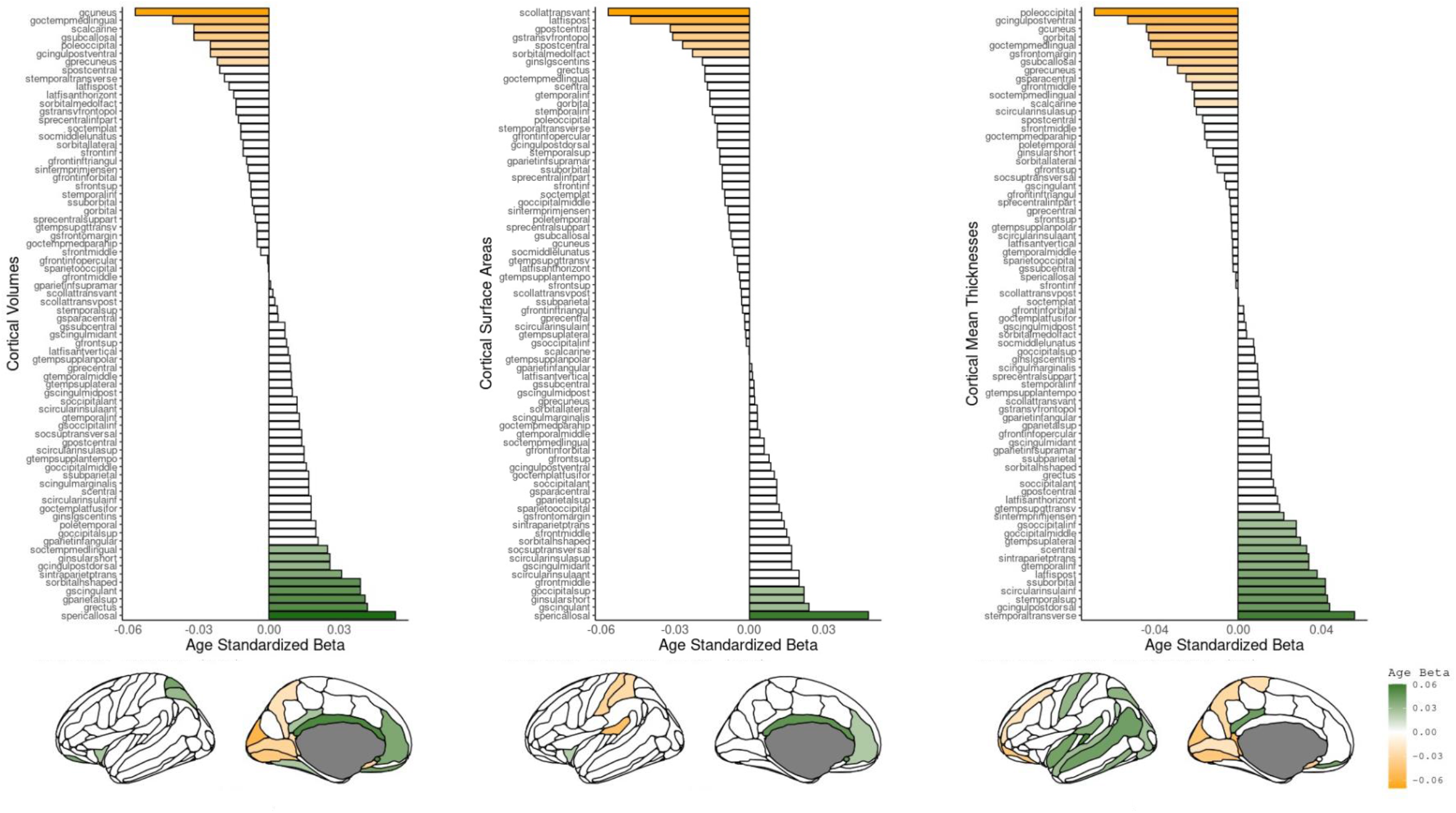
Linear Age Differences in Asymmetry Index (AI) across Cortical Volumes, Surface Areas, and Mean Thicknesses. Positive linear age-standardized beta indicates an increase in AI with linear (green) and a negative slope indicates a decrease in AI with linear age (yellow). Colors indicate significance at 0.05/(5*306). Figures made with ggseg (Mowinckel & Vidal-Piñeiro, 2019).

#### 3. Subcortical Regions

The L-R caudate, the accumbens area, and the thalamus was greater in females compared to males (Figure 3). All hippocampal subfields had an AI that was greater in females compared to males, whereas 48% (12/25) of thalamic subfields were larger in females and 16% (3/25) in males. The L-R pallidum and the amygdala was greater in males compared to females (Figure 3). All amygdala subfields had a larger AI in males compared to females. The L-R thalamus and ventral DC increased with age, whereas the L-R caudate, the pallidum, the accumbens area, and the hippocampus decreased with age (Figure 3). The L-R thalamus also increased with quadratic age.

**Figure 3.**
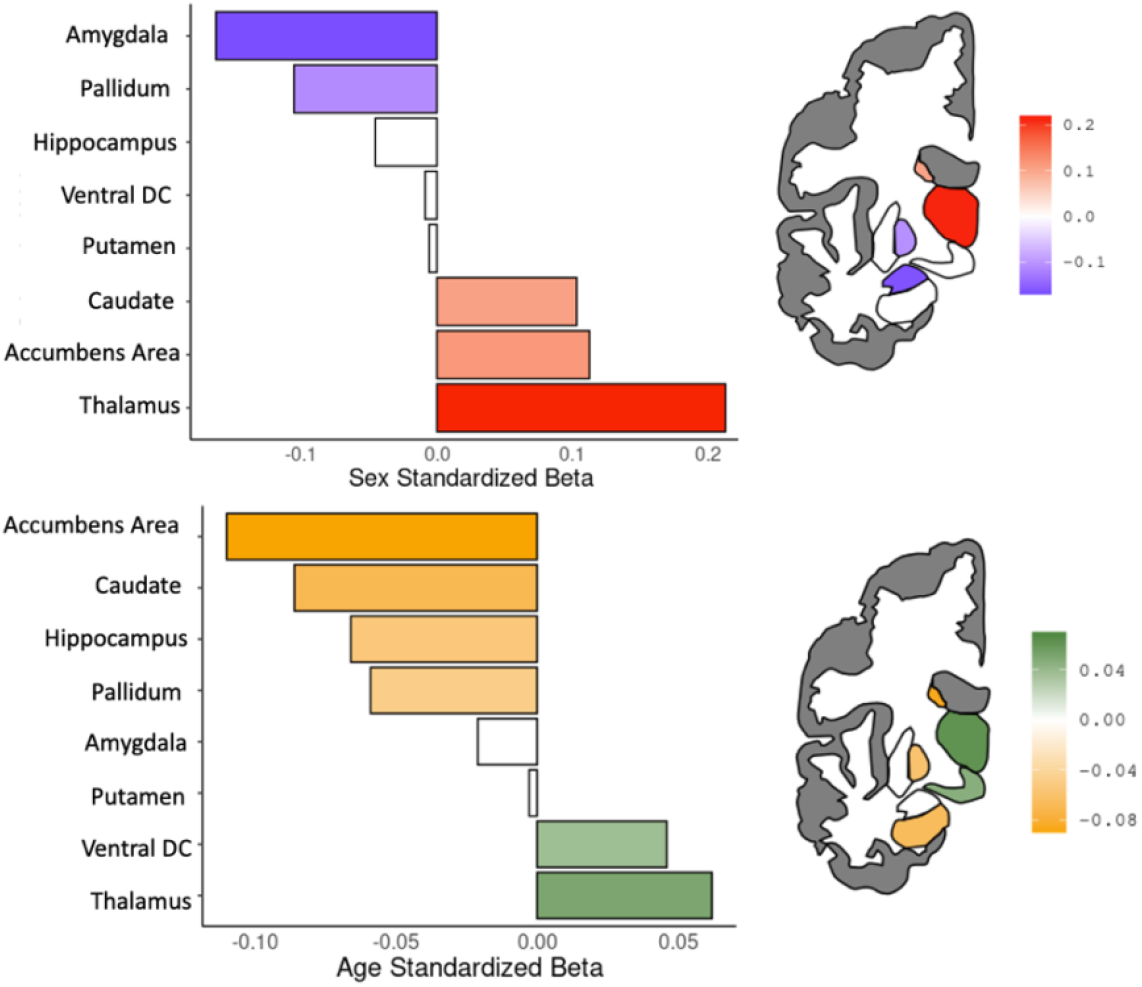
Sex and Linear Age Differences in Asymmetry Index (AI) across Subcortical Volumes. Colors indicate significance at 0.05/(5*306). Positive sex standardized betas indicate that females have a greater AI than males (red) and a negative slope indicates that males have a greater AI than females (blue). Positive linear age-standardized beta indicates an increase in AI with linear (green) and a negative slope indicates a decrease in AI with linear age (yellow). DC: Diencephalon. Figures made with ggseg (Mowinckel & Vidal-Piñeiro, 2019).

There was a significant interaction of sex with linear age in the pallidum (β = 0.02), the cortical nucleus of the amygdala (β = -0.05), the CA3 hippocampal head (β = -0.06), and the thalamic lateral geniculate (β = -0.06), ventral posterolateral (β = -0.06), mediodorsal medial magnocellular (β = -0.05), and ventral medial (β = -0.06) nuclei.

The AIs of regions with sex differences went in the same direction for both sexes except for two thalamic nucleus volumes. Females had a leftward asymmetry, whereas males had a rightward asymmetry in the ventral medial and ventral posterolateral thalamic nuclei volumes (β = 0.09 and β = 0.26, respectively; Supplemental Tables SG1).

#### 4. Cerebellar Grey Matter Volumes

The L-R crus I lobule (β = 0.15) was greater in females, whereas the L-R VIIA, VIIIA, and X lobules were larger in males (β = -0.12, β = -0.17, β = -0.06, respectively). The L-R VI, VIIB, and VIIIA lobules decreased with age (β = -0.03, β = -0.04, β = -0.04, respectively) and the crus II, VIIIB, and X lobules increased with age (β = 0.04, β = 0.03, β = 0.04, respectively). The L-R VIIB and crus II lobules had a negative quadratic age effect (β = -0.03, for both) and there was a significant interaction of linear age with sex in the AIs of the crus II and X lobules (β = -0.05, β = -0.06, respectively). Males had a leftward asymmetry and females a rightward asymmetry in the VIIIB lobule (β = -0.17).

#### 5. Handedness

Left-handed individuals had greater rightward asymmetries in TSA, Cerebral WMV, and Cerebral GMV, whereas they had greater leftward asymmetries in Total MCT and Cortical GMV. Handedness also affected 4 regional volumes, 6 cortical mean thicknesses, and 3 cortical surface areas. Left-handed individuals had a greater rightward asymmetry in the lateral posterior thalamic nucleus volume and a greater leftward asymmetry in the cerebellar VIIIB lobule, the ventral anterior thalamic nucleus, and the orbital gyrus volumes. In terms of cortical mean thicknesses, left-handed individuals had a greater rightward asymmetry in the postcentral sulcus and a greater leftward asymmetry in the suborbital sulcus, the pericallosal sulcus, the medial orbital sulcus, the gyrus rectus, and the anterior part of the cingulate gyrus, and sulcus (ACC). The surface area of the ACC also differed between right- and left-handed individuals. However, left-handed individuals had a greater rightward asymmetry in the surface areas of the ACC and the middle-posterior part of the cingulate gyrus and sulcus and they had a greater leftward asymmetry in the orbital sulci compared to right-handed individuals. Finally, the asymmetries of right-handed and ambidextrous individuals did not differ across global and regional measures.

## 4. Discussion

The present study reports the brain measures to take into consideration when examining group differences in cerebral asymmetries that are independent of brain size. We found that the L+R Measure, the TCM, and the L-R TCM predict the AI of over 89% of investigated regions. The relationship between these predictors and the AIs deviated little from linearity to warrant the use of non-linear models. We used the case of sex differences in brain asymmetries to illustrate the consequences of omitting these predictors when examining group differences in asymmetries. We found that removing either the L+R Measure, the TCM, or the L-R TCM changed the magnitude and significance of the sex differences in asymmetry. Finally, we reported the effects of sex, age, and their interactions when including the L+R Measure, the TCM, and the L-R TCM, which, based on our results, should be considered by future studies examining group differences in brain asymmetries.

### L+R Measure

In contrast to popular belief (Guadalupe et al., 2017; X.-Z. Kong et al., 2020; Kurth et al., 2015), we found that dividing the L-R Measure by the L+R Measure does not ensure that the index does not simply scale with brain size. Dividing the L-R Measure would sufficiently adjust for the L+R Measure only if the intercept of their relationship was null. However, the intercept is not null and the L+R Measure was a significant predictor of the AI. Omitting the L+R Measure significantly changed the magnitude and the significance of the sex differences in 42% of regions. Thus, researchers should include the L+R Measure as a covariate in their analyses of group differences in asymmetry.

If the L+R Measure is included as a covariate in their model, there is no need to divide the L-R Measure by its L+R Measure. One could examine brain asymmetries by analyzing L-R Measures with the appropriate covariates. However, the standard AI is convenient since it has the same scale across regions of different sizes. For this reason, we recommend using the standard AI as long as all appropriate covariates are included.

### Global Brain Size

Our findings corroborate the current practice of adjusting for brain size (e.g., TBV) to report group differences in cerebral asymmetry that are independent of individual differences in global brain size. In line with previous studies (Guadalupe et al., 2017; Kang et al., 2015; X.-Z. Kong et al., 2018), TCM was a significant predictor of a majority of regions, even when including the L+R Measure in the model. Although Kong and colleagues (2018) found an effect of ICV on the AI of overall cortical thickness but not on the AI of overall surface area, we found that TBV predicted both overall surface area and mean thickness, with greater asymmetries in larger brains.

While omitting TCM influenced the magnitude of the sex effect in volumes and surface areas, it did not influence the magnitude of the sex differences in AIs across cortical mean thicknesses. This may be because mean thickness is quite small, allowing for little variability across individuals. Removing total MCT nonetheless influenced the significance of the sex effects: four regions that were significant in the model with all the covariates were no longer significant when removing total MCT and one significant region in the model without total MCT was no longer significant in the model with all the covariates. Therefore, we suggest that Total MCT be maintained as a covariate when investigating sex differences in asymmetries. Finally, since Total MCT is a significant predictor of asymmetry across 97% of mean thicknesses, omitting Total MCT could influence the magnitude and significance of other grouping variables of interest and should be taken into account by future studies.

### L-R TCM

To our knowledge, this paper is the first to examine the effects of global brain asymmetry (L-R TCM) on group differences in regional asymmetries. We found that the asymmetry between the left and the right TCMs predicts the AI of a majority of regions (85%). Removing the L-R TCM changed the magnitude of the reported sex differences in asymmetry across volumes and mean thicknesses but not surface areas. However, removing the L-R TSA led to changes in the significance in 8 sex differences in asymmetry. Therefore, we recommend that the L-R TCM still be considered when investigating sex differences in surface areas to disentangle global from local differences in asymmetry. Finally, since the L-R TSA is a significant predictor of the asymmetry in 81% of surface areas, omitting the L-R TSA could influence the magnitude and significance of other grouping variables of interest and so should be considered as a covariate by future studies.

These analyses lead us to recommend that studies of brain asymmetry systematically adjust on all three covariates: L+R Measure, TCM, and L-R TCM. In essence, this amounts to studying residual asymmetries: asymmetries once global brain size and asymmetry factors are taken into account. Some investigators may feel that absolute asymmetries are more relevant for their purposes, or perhaps more functionally significant. They may be. The problem is that when group differences in absolute regional AI are found, one does not know whether those differences are specific to this region, or whether they are due to differences in region size, brain size, or global brain asymmetry. By comparing models with and without these covariates, researchers will assess the extent to which observed effects are regional, and the extent to which they are global, allowing for a more fine-grained understanding of variations in brain asymmetries. We applied this approach to a systematic study of the relation between residual cerebral asymmetries and sex, age and handedness in the UK Biobank, with the following results.

### Sex Effects

Asymmetries in males and females were generally in the same direction and the majority of sex differences reflected a more pronounced asymmetry pattern in one sex compared to the other. Consistent with Kong and colleagues (2018) and Koelkebeck and colleagues (2014), we found that males have a greater rightward asymmetry in overall surface area compared to females, beyond that expected from the differences in brain size. In line with Kong and colleagues (2018), we found a more pronounced leftward asymmetry in males compared to females in the surface areas of the superior temporal and supramarginal gyri in the Desikan-Killiany-Trouville atlas and supramarginal and lateral aspect of the superior temporal gyri in the Destrieux atlas. We additionally reported a greater rightward asymmetry in females compared to males in the inferior parietal gyrus of the DKT atlas and the angular gyrus of the Destrieux atlas. We did not find a sex difference in asymmetry in the mean thickness of the enthorinal and parahippocampal mean thicknesses as indicated by Kong and colleagues (2018), nor did we find a sex difference in the putamen as suggested by Guadalupe and colleagues (2017). These different results between our study and the literature can be explained by differences in raw AIs (e.g., putamen) or differences between samples (e.g., sample age).

With such a large dataset, we reported more significant sex differences in asymmetries than in previous studies (Guadalupe et al., 2015; Koelkebeck et al., 2014; X.-Z. Kong et al., 2018, 2020): We found 55 regions with greater asymmetries in women and 67 with greater asymmetries in males. Although sex differences in asymmetries were generally consistent across segmentation algorithms and models, there were some exceptions: We observed a greater leftward asymmetry in males in the superior frontal gyrus surface area in the Destrieux atlas but not with the Desikan-Killiany-Trouville atlas. We also only reported sex differences in the asymmetry of mean thicknesses in the superior temporal gyrus, rostral middle frontal gyrus, and the paracentral gyrus when the L-R TCM and L+R Measure were included in the models. These results further highlight the importance of adjusting for these brain measures to report sex differences in regional asymmetries that are independent of brain size and global asymmetry.

Several regions with a greater rightward asymmetry in males across surface areas had a greater leftward asymmetry in females across mean thicknesses. The presence of sex differences in cortical thickness asymmetry across regions of the language network, such as the supramarginal gyrus, the temporal transverse sulcus, planum polare of the superior temporal gyrus, and the anterior transverse temporal gyrus of Heschl, contrasts with Kong and colleagues’ (2018) findings, who did not report sex differences in these regions and, in turn, hypothesized that sex differences in the asymmetry of cortical thicknesses were independent of sex differences in performance on language tasks. Further structural and functional studies are needed to evaluate whether sex differences in linguistic skills are associated with sex differences in asymmetries across the volumes, surface areas, and/or cortical thicknesses of regions in the language network.

### Age Effects

Age did not influence brain asymmetries across a majority of regions in the UK Biobank. Although our sample is much older than Kong and colleagues (2018), we found the greater leftward asymmetry with increased age in the mean thickness of the superior temporal gyrus. Therefore, age effects on mean thickness asymmetries in certain regions may remain stable across one’s lifetime. In contrast with Kong and colleagues’ (2018) study, we did not find a greater leftward asymmetry with increasing age in the surface area of the entorhinal cortex. Instead, we found a greater leftward asymmetry with increasing age in the surface areas and volumes of the cingulate regions and rightward asymmetry with increasing age in the surface areas and volumes of the postcentral gyrus and sulcus and the posterior ramus of the lateral sulcus. Considering that we are (to our knowledge) the only ones with Kong and colleagues (2018) to report age effects across surface area asymmetries, our findings must be replicated to be judged as robust.

### Handedness

Our results further support that handedness – one of the most evident functional lateralization patterns (Papadatou-Pastou et al., 2020) – was not associated with the majority of investigated asymmetries in volumes, mean thicknesses, and surface areas. Although our findings coincide with the latest large-scale studies examining brain lateralization with similar asymmetry indices (Guadalupe et al., 2017; X.-Z. Kong et al., 2018), we similarly found an effect of handedness in the surface areas of the anterior medial cingulate regions and the mean thicknesses of the postcentral region (Sha et al., 2021). Cerebral asymmetries may nonetheless more systematically vary as a function of handedness when measured by other indices. For instance, a recent study using an automated registration-based approach reported that the horizontal and vertical skew (L-R differences along the anterior-posterior and dorsal-ventral axes, respectively) differed between left- and right-handed individuals in the UK Biobank and Human Connectomes Project (X.-Z. Kong et al., 2018).

### Limitations

The UK Biobank dataset consists of older adults that are healthier and have a higher socioeconomic status than the UK population (Fry et al., 2017). The UK Biobank imaging sample also shows a healthy bias compared to the UK Biobank sample (Lyall et al., 2021). Therefore, future studies should consider that the associations between the asymmetry markers we generated and mental health disorders are likely not generalizable to the UK and UK Biobank population, as studies with the UK Biobank imaging sample may underestimate exposure and outcome estimates (Lyall et al., 2021). The present study nonetheless corresponds to the largest study of sex and age effects and interactions on brain asymmetries. We appropriately controlled for individual differences in brain size and used data from a few imaging sites with similar MRI scanner characteristics and acquisition sequences, reducing the likelihood of obtaining biased estimates from different imaging parameters.

## Conclusion

We find that the widely-used AI significantly scales with brain size. Using the case of sex differences, we illustrate that the L+R Measure, the TCM, and the L-R TCM should be considered to report unbiased group differences in cerebral asymmetries. Taking into account the appropriate covariates will not only shed light on debated brain asymmetries across healthy controls but also on the variations in brain asymmetry associated with differences in cognition and mental health. Although numerous studies examining the link between mental health disorders and asymmetry find that, given the small effect sizes, structural brain asymmetry alone is unlikely to be a useful biomarker of many disorders for individual-level prediction or diagnosis (X.-Z. Kong et al., 2020; Postema et al., 2021) studying brain asymmetry may contribute to understanding the neurobiological underpinnings of cognition and psychiatric disorders.

## Abbreviations

L: Left Hemisphere
R: Right Hemisphere
AI: Asymmetry Index = (L-R)/(L+R)
TBV: Total Brain Volume
TSA: Total Surface Area
Total MCT: Total Mean Cortical Thickness
TCM: Total Cerebral Measure (i.e., TBV or TSA or Total MCT)
Measure: Volume, Mean Thickness, Surface Area
L+R: L+R Measure
L-R: L-R Measure
L-R TCM: L-R Total Cerebral Measure (i.e., L-R TBV or TSA or Total MCT)

## Acknowledgments

This work received support under the program “Investissements d’Avenir” launched by the French Government and implemented by l’Agence Nationale de la recherche (ANR) with the references ANR-17-EURE-0017 and ANR-10-IDEX-0001-02 PSL. Funding was also obtained from Fondation pour l’Audition (FPA RD-2016-8 research grant). This research has been conducted using the UK Biobank Resource.

## Footnotes

1 Consider Left (L) and Right (R) measures for a given region. L is regressed on R, with a>0: L=aR+b (error term removed for simplicity). If L and R are correlated and a<1 then L-R= (a-1)R+b will be negatively correlated with R and dividing by L+R will not remove this correlation: AI=(L-R)/(L+R)= ((a-1)R+b)/((a+1)R+b). Therefore, the nature of the relationship with R depends on the values of a and b. Since the AI is correlated with R in most cases, it will also be correlated with L+R.

2 We could not use the R2 or an F-test because the R2 is not valid for non-linear models (Spiess & Neumeyer, 2010) and the F-test is only for nested models. Finally, we could not compare the AIC or BIC across models because splines and linear models use different likelihood functions.

