## Supplemental Information for "Comparing Brain Asymmetries Independently of Brain Size"

### Table of Contents


***1 – Total Brain Volume (TBV) Choice*** ***2***

***2 - Scanner Site*** ***6***

***3 - Handedness*** ***6***

***4- Regions with a Residual Standard Error (SE) greater than 1*** ***7***

***5- Sex Differences*** ***7***

***5- Exploratory analyses*** ***9***

**1.** 1

**2.** 3

**3.** 5

**4.** 6

a) 6
b) 6
c) 1
d) 2

**5.** 3

a. 3
b) 5
c) 1

**6) Comparing Subcortical AIs and Sex and Age Effects** **24**

a) 1
b) 1

***6 - R packages*** ***25***

### 1 – Total Brain Volume (TBV) Choice

This section was taken from the Williams and colleagues (2021) supplemental information: https://osf.io/x5egr/ and included in these supplements to facilitate access.

Both TBV and Total Intracranial Volume (TIV) have been used to adjust for brain allometry to avoid a bias toward isometry when estimating allometric scaling coefficients of large regions (e.g., Jong et al., 2017; Peyre et al., 2020; Reardon et al., 2016; Williams et al., 2021). However, since segmentations yielding the highest number of regions were favored and because we are interested in examining how much a particular volume contributes to the total brain volume at the time of the study (and not at maximal lifetime total brain volume; Herbert et al., 2003; O'Brien et al., 2006), TBV was chosen over TIV. The correlation between TBV and TIV was 88.93%.

We did not use the TBV provided by ASEG (26515) due to a mismatch for some participants between the provided TBV measure (26515) and the one calculated by summing the global measures provided by ASEG (*Calculated ASEG TBV*): Total GMV Volume (26518) + Cerebellum WMV (Left: 26556, Right: 26587) + Cerebral WMV (Left: 26553, Right: 26584). This mismatch yielded a correlation of 99.73% between TBV measures (Figure S1). We used the *Calculated ASEG TBV* as our measure of TBV since it correlated at 99.82% with the TBV we calculated from summing the regions of interest in the present study (*Calculated TBV*, Figure S2). This allowed us to obtain a TBV measure for a maximum of participants (N = 40 028). (We would otherwise, for instance, have to exclude all individuals that do not have the FAST cerebellum segmentations even though they have all the other segmentations).

As for the Left and Right TBV measures, we used the following ASEG segmentations: Cerebellum GMV (Left: 26557, Right: 26588), Cerebral GMV (Left: 26552, Right: 26583), Cerebellum WMV (Left: 26556, Right: 26587), Cerebral WMV (Left: 26553, Right: 26584), and the total subcortical volume we calculated with the ASEG and Freesurfer subsegmentations. The Left and Right TBV measures from a majority of ASEG segmentations (*Left +Right ASEG TBV*) was favored over calculating the Left and Right TBV measures from the sum of all left and right brain volumes of interest (*Calculated TBV)*, to obtain a Left and Right TBV measure for a maximum of participants (N = 38 710, instead of N = 37 945). *The Left +Right ASEG TBV* Measure and the *Calculated TBV* correlated at 99.80% (Figure S3).

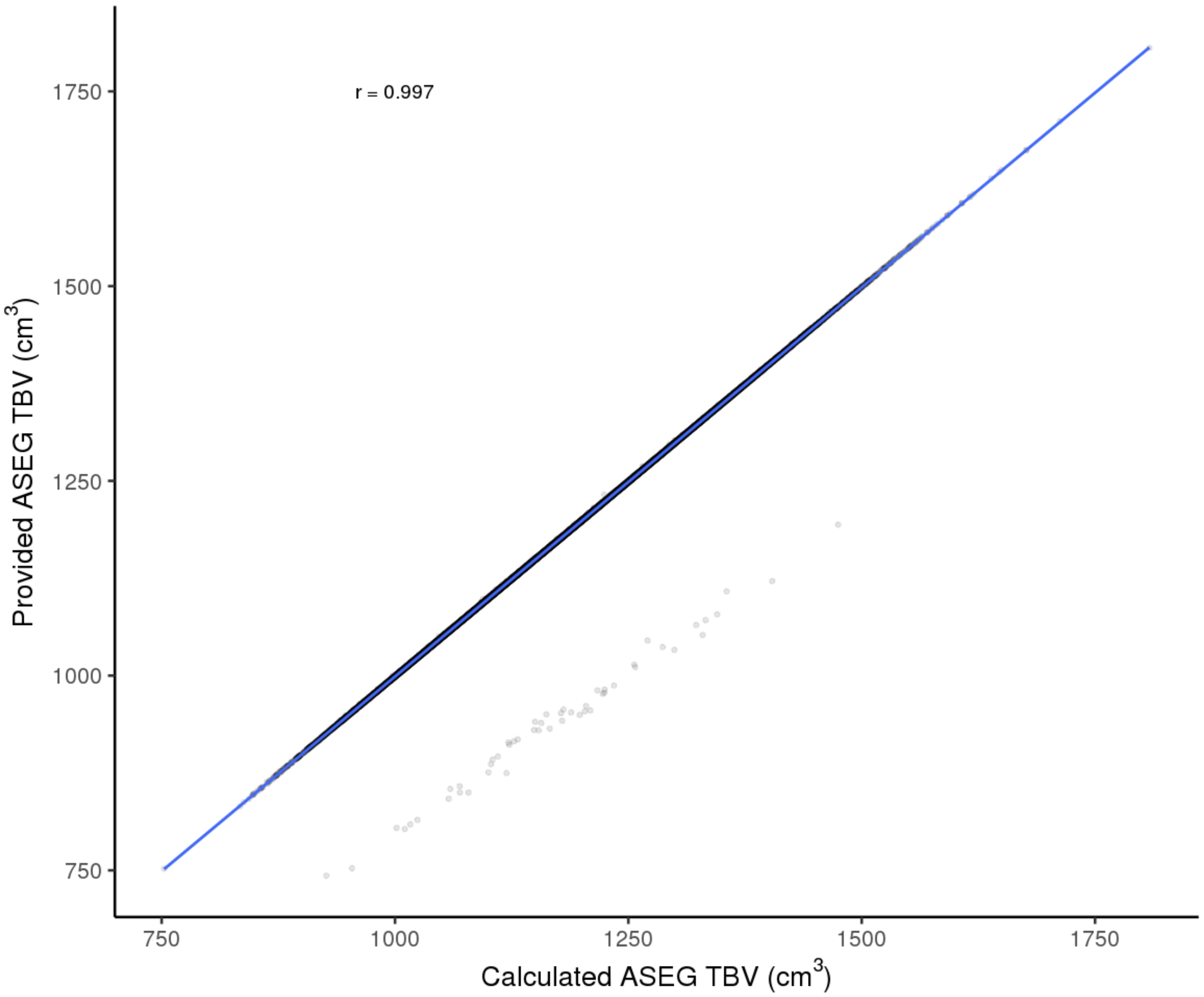


***Figure S1*. Correlation between the Provided ASEG Total Brain Volume (TBV) Measure and the Calculated Sum of Global Measures from the Freesurfer ASEG Segmentations.**

The provided TBV corresponds to data-field 26515 and the Calculated ASEG TBV to the sum of Total Grey Matter Volume (data-field 26518), Cerebellum White Matter Volume (Left: data-field 26556, Right: data-field 26587), Cerebral White Matter Volume (Left: data-field 26553, Right: data-field 26584). r, rho = 99.73%.

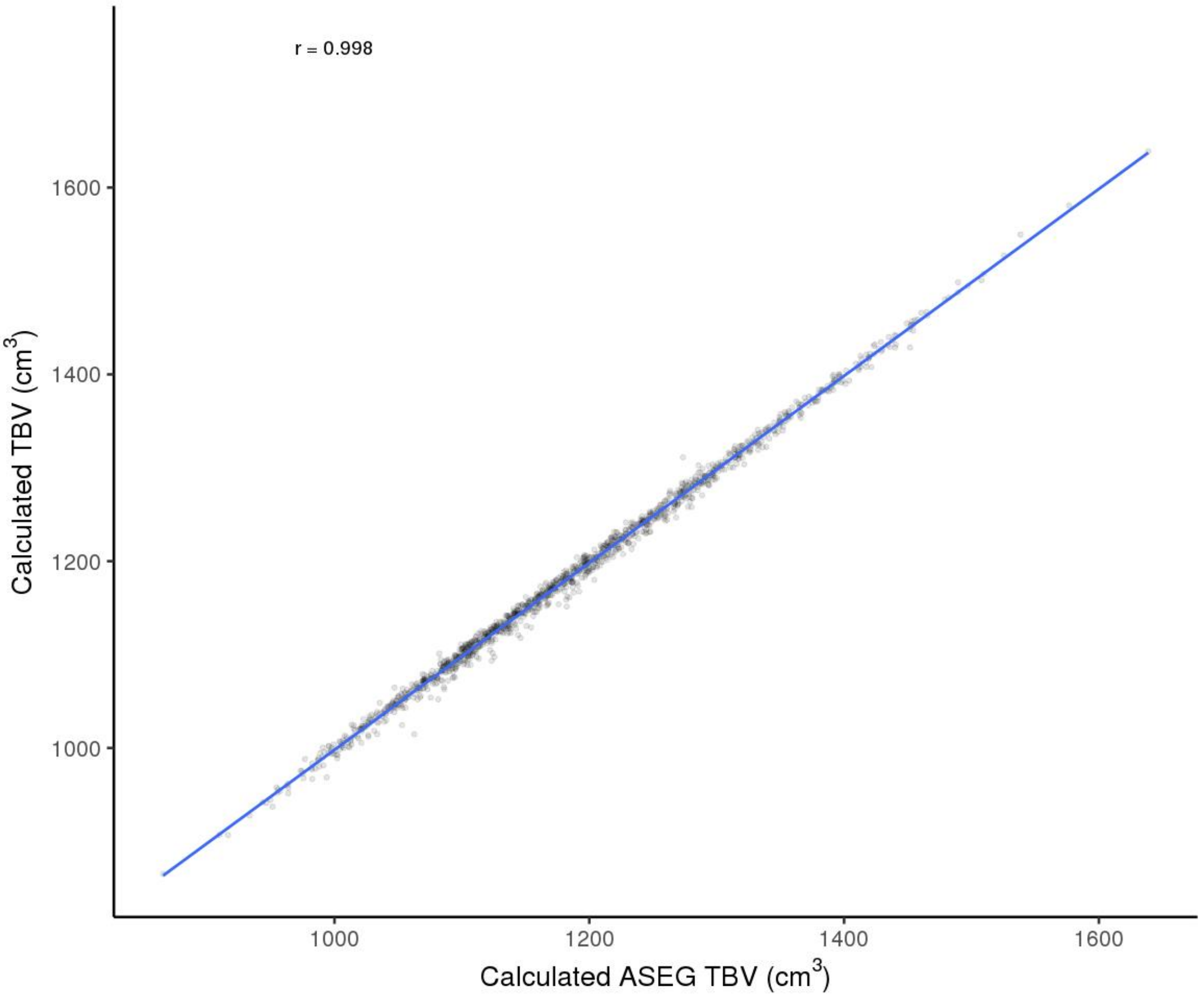


***Figure S2*. Correlation between the Calculated ASEG Total Brain Volume (TBV) Measure and the Calculated TBV from the Regions examined in this study.**

The Calculated ASEG TBV corresponds to the sum of Total Grey Matter Volume (data-field 26518), Cerebellum White Matter Volume (Left: data-field 26556, Right: data-field 26587), Cerebral White Matter Volume (Left: data-field 26553, Right: data-field 26584). r, Pearson r.

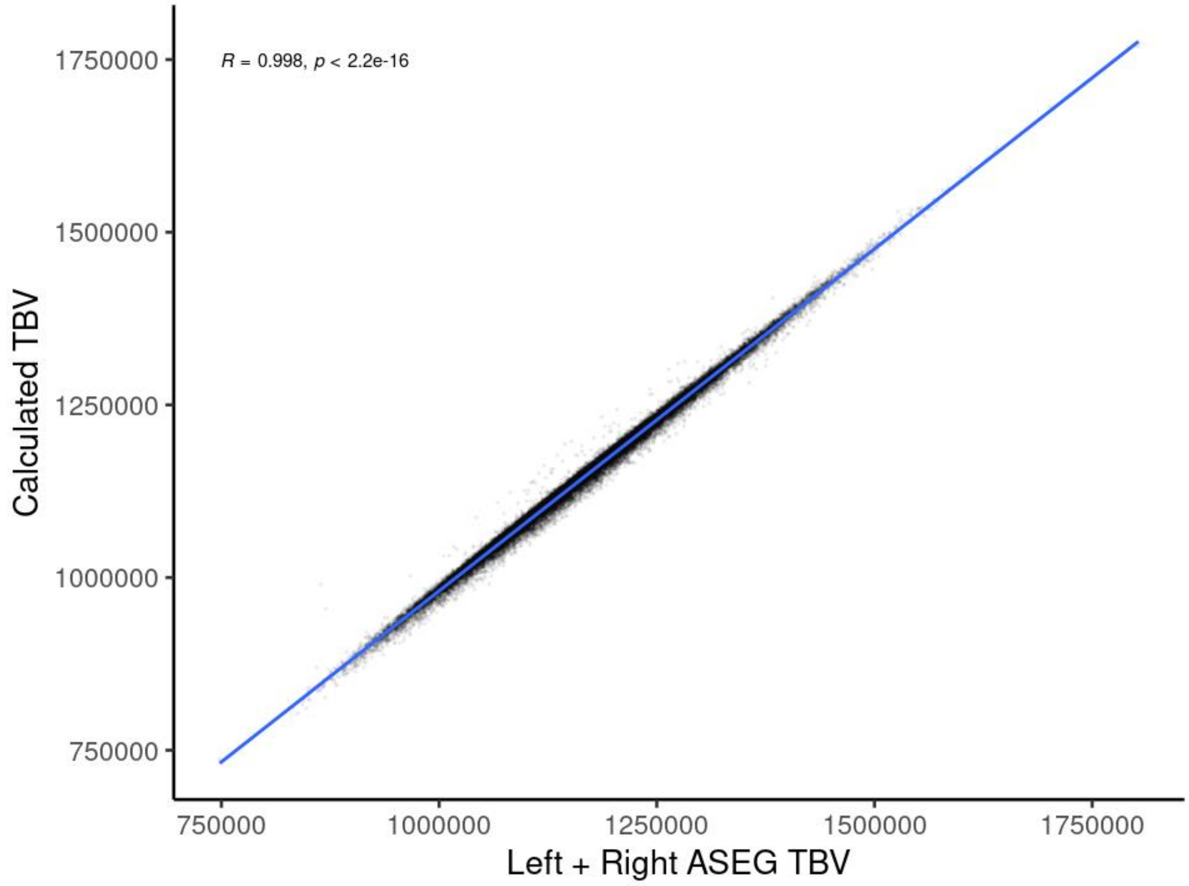


***Figure S3*. Correlation between the Calculated Left + Right ASEG Total Brain Volume (TBV) Measure and the Calculated TBV from the Regions examined in this study.** The Calculated ASEG TBV corresponds to the sum of Total Grey Matter Volume (data-field 26518), Cerebellum White Matter Volume (Left: data-field 26556, Right: data-field 26587), Cerebral White Matter Volume (Left: data-field 26553, Right: data-field 26584). r, Pearson r.

### 2 - Scanner Site

The number of participants differed across sites ($\chi 2$ (2, N = 40 028) = 16 107, $p < 2.2e-16$) and between sexes ($\chi 2$ (2, N = 40 028) = 27.1, p =1.314e-06), with more female than male participants across sites (Site 11025: Male N = 12033 and Female N = 12938, Site 11026: Male N = 2318 and Female N = 2752, and Site 11027: Male N = 4535 and Female N = 5452). Age differed between Site 11025 and Site 11026 (b = 2.27 years, $t$ (40025) = 19.63, $p < 2.2e-16$), between Site 11025 and Site 11027 ($b$ = 1.40 years, $t$ (40025) = 15.78, $p < 2.2e-16$), and between Site 11026 and Site 11027 ($b$ = -0.87 years, $t$ (40025) = -6.70, p = 2.1e-11).

### 3 - Handedness

The number of participants differed by handedness ($\chi 2$ (2, N = 39 854) = 56 584, $p < 2.2e-16$) and between sexes ($\chi 2$ (2, N = 39 854) = 69.1, p =9.902e-16), Table S1. Age differed between Right- and Left-handed individuals (b = -0.36 years, t (40025) = -2.75, SE = 0.13, p = $6.00 \times 10^{-3}$) and between Right-handed and Ambidextrous individuals ($b$ = 1.59 years, $t$ (40025) = 4.90, p = $9.78 \times 10^{-7}$).

Table S1. Handedness as a function of sex

| Handedness | Female, N | Male, N |
|---|---|---|
| Left-handed | 1766 | 1945 |
| Right-handed | 19065 | 16531 |
| Use both right and left hands equally | 236 | 311 |
| Missing | 75 | 99 |

#### 4- Regions with a Residual Standard Error (SE) greater than 1

We conducted exploratory analyses to identify the cerebral predictors responsible for the reduction in RSE in the six regions that had a reduction in RSE that was larger than 1%. We ran the three models for each region. Each model consisted of the cerebral AI as a function of either the L+R Measure, the TCM, or the L-R TCM. We found that the reduction in RSE above 1% was due to the non-linear relationship between the AI and the L+R Measure across regional models and the non-linear relationship between the AI and the Total Cerebral AI for global measures (Supplemental Tables C6 and Files S1-3).

We conducted additional analyses in the regions with a reduction in RSE that was greater than 1% in Q2. For each of these regions, we selected the model with the smallest RSE and identified which predictor(s) had a non-linear relationship with the AI to create a new non-linear model for the AI in question. We then conducted a Z-test between the sex coefficient of the linear model and the new non-linear model to examine whether omitting non-linearity influenced reported sex differences. We found that omitting the non-linear relationship in these regions did not influence the magnitude of the reported sex difference ($p < 0.05$; Supplemental File S4).

### 5- Sex Differences

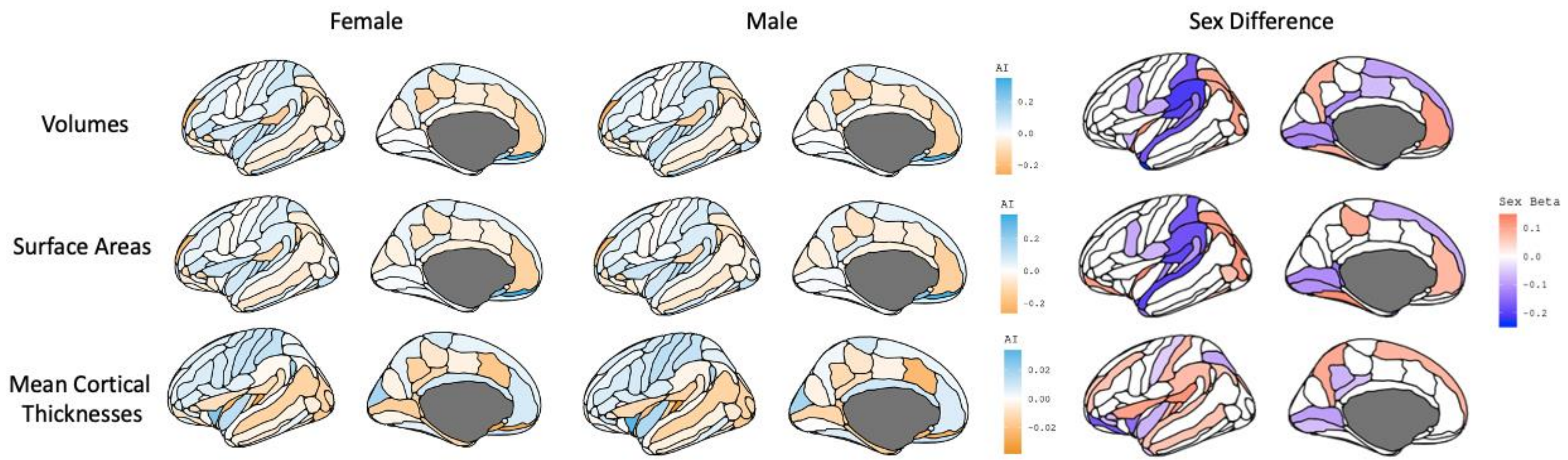


***Figure S1***. *Asymmetry Index (AI) by Sex and Sex Differences in AIs across Cortical Volumes, Surface Areas, and Mean Thicknesses.* Light Blue Regions: Left > Right Measure and Orange Regions: Right > Left Measure. Red Regions: Positive Sex Standardized Beta indicating that Female > Male AI and Dark Blue Regions: Negative Sex Standardized Beta indicating that Male > Female AI. White regions: Sex difference is not significant at 0.05/(5*306). Figures made with ggseg.

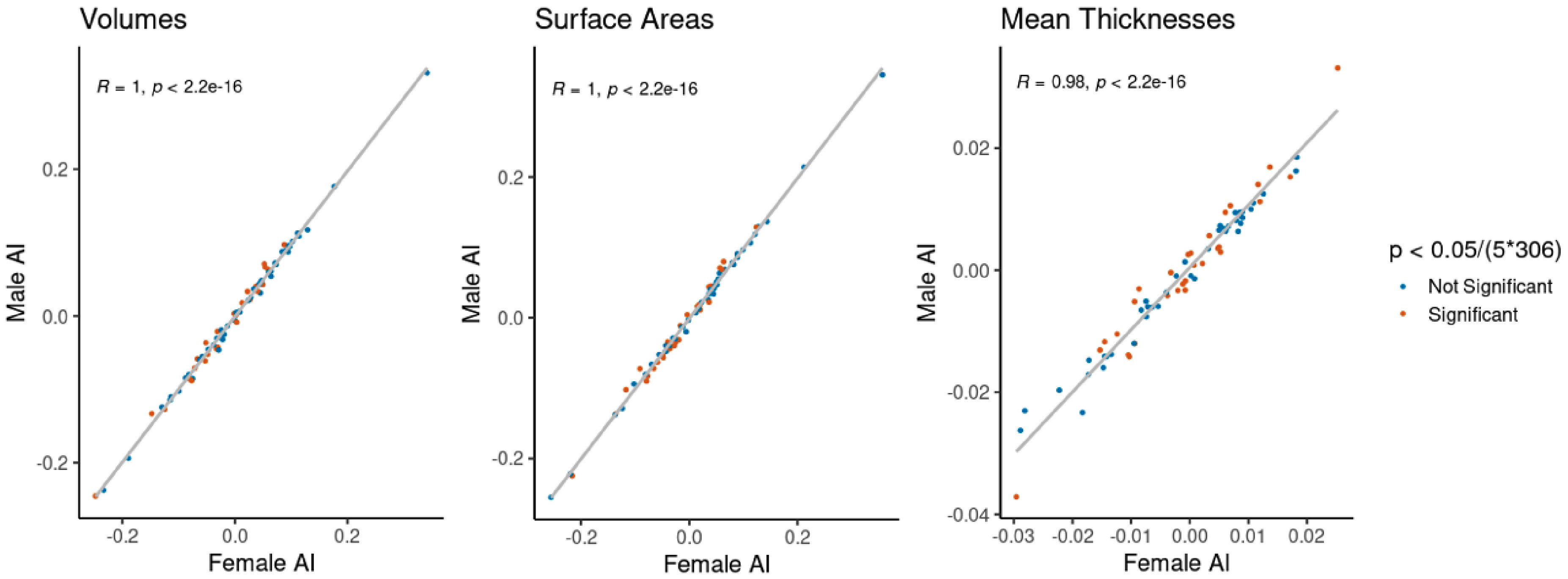


***Figure S2****. Correlation of the Asymmetry Index (AI) by Sex across Cortical Volumes, Surface Areas, and Mean Thicknesses*. Regions with significant sex differences are in orange and nonsignificant in blue.

### 5- Exploratory analyses

The following analyses were not preregistered unless indicated by an *.

#### 1. Global Asymmetry Markers

We derived three global measures of asymmetry to quantify a person's level of absolute asymmetry across either all volumes, surface areas, or mean thicknesses (equation 13). When regions were embedded in each other, we included the smallest measures of brain asymmetries (e.g., we used the AI of each hippocampal subfield instead of the AI of the whole hippocampus). We then examined sex differences in the 3 global asymmetry markers (volume, surface, thickness). The significance threshold was set to 0.05/(5*306).

**Equation 13**: $\text{Global Asymmetry Marker} = \sqrt{(\sum(\text{Regional Asymmetry Indices})^2) / N}$

The Global Asymmetry Marker was larger in males compared to females for volumes ($\beta$ = -0.13) and surface areas ($\beta$ = -0.08), and was larger in females for mean thicknesses ($\beta$ = 0.11; Supplemental Information Figure S3). The Global Asymmetry Marker increased with linear and quadratic age for volumes ($\beta$ = 0.10 and $\beta$ = 0.02, respectively) and mean thicknesses ($\beta$ = 0.09 and $\beta$ = 0.04, respectively). The L-R TCM negatively predicted the Global Mean Thickness Asymmetry Marker ($\beta$ = -0.05) and Total MCT negatively predicted the Global Mean Thickness Asymmetry Marker ($\beta$ = -0.24; Descriptive Statistics, Supplemental Table A9 and Results, Supplemental Table F8). There were no sex differences in variance across Global Asymmetry Markers (Supplemental Table H5).

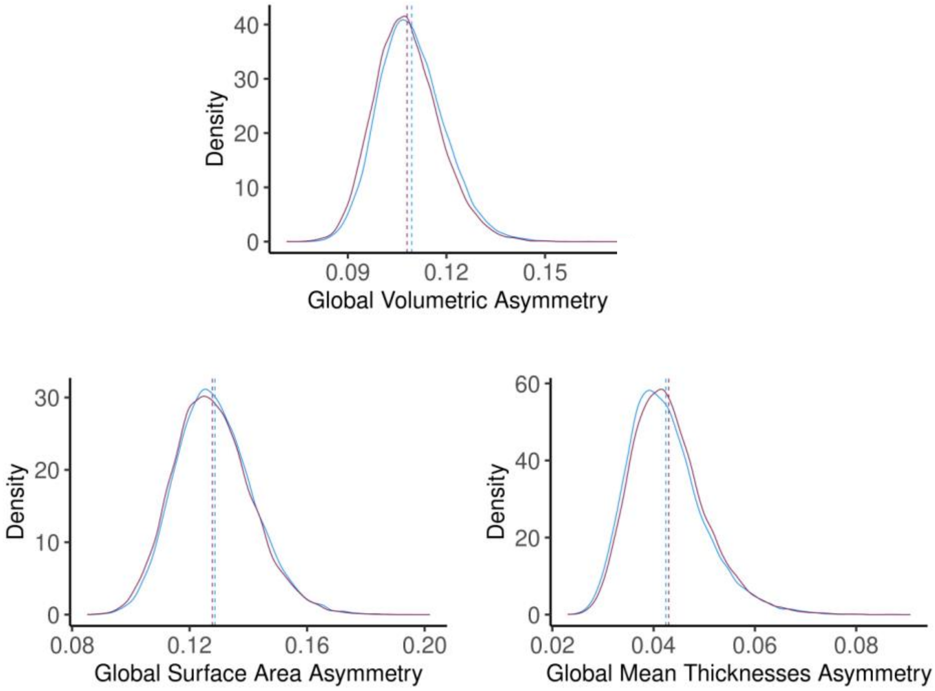


***Figure S3.*** *Sex Differences in Global Asymmetry Markers across Volumes, Cortical Surface Areas, Cortical Mean Thicknesses.* Mean (dashed lines) differences were found across measures (violet: female, blue: male). The Global Asymmetry Marker corresponds to the square root of the sum of squared asymmetry indices divided by the number of regions for that measure (148 for volumes and 74 for mean thicknesses and surface areas). Significance set to 0.05/(306*5).

### 2. Global Asymmetry Deviance Markers*

As done in a previous paper (Williams et al., 2021), we created person-level deviance markers of asymmetry by extracting the residuals of equation 12 for each region. These markers reflect the extent to which a person deviates from their predicted asymmetry in a region given their age, sex, L+R Measure, TCM, and L-R TCM.

Finally, with the person-level deviance markers of asymmetry, we generated three global deviance markers of asymmetry: one for volumes, one for surface areas, and one for mean thicknesses (equation 14). These global markers reflect the extent to which a person deviates from their predicted asymmetry given their age, sex, L+R Measure, TCM, and L-R TCM across all volumes, mean thicknesses, or surface areas. We then performed a t-test to examine sex differences on these markers. The significance threshold was set to 0.05/(5*306).

**Equation 14**: Global Asymmetry Deviance Marker = $\sqrt{}$ ($\sum$ (Person-Level Asymmetry Deviance Markers)$^2$) / N

Males had larger global asymmetry deviance markers compared to females across volumes ($t(37673) = 17.97$, $p < 2.2e\text{-}16$, $d = 0.18$) and surface areas ($t(37790) = 10.11$, $p < 2.2e\text{-}16$, $d = 0.10$), while females had larger global asymmetry deviance markers compared to males across mean thicknesses ($t(38534) = 11.62$, $p < 2.2e\text{-}16$, $d = -0.12$; Supplemental Information Figure S3; Descriptive Statistics, Supplemental Table A10).

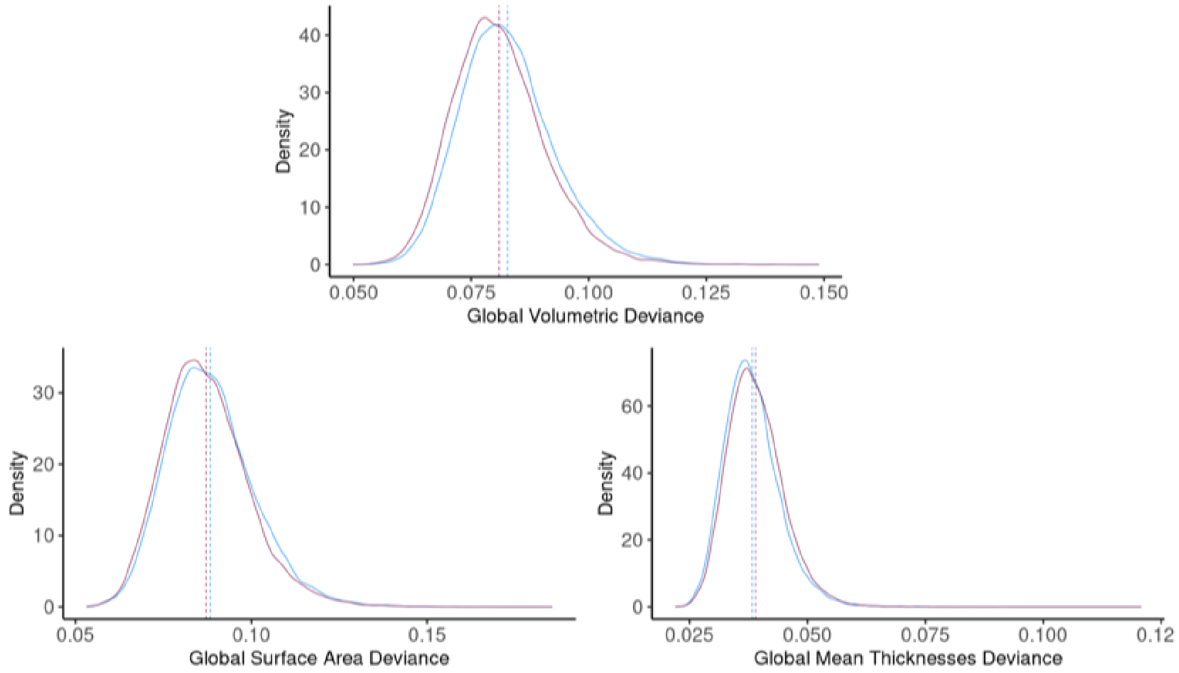


***Figure S4.*** *Sex Differences in Global Asymmetry Deviances across Volumes, Cortical Surface Areas, Cortical Mean Thicknesses.* Mean (dashed lines) differences were found across measures (violet: female, blue: male). Global Asymmetry Model corresponds to the square root of the sum of squared asymmetry indices divided by the number of regions for that measure (148 for volumes and 74 for mean thicknesses and surface areas). Significance set to 0.05/(306*5).

#### 3. Regional Sex Differences in Variance*

As preregistered, we examined sex differences in asymmetry variance. We ran the equations from Q4 with continuous variables that are centred but not divided by 1SD and extracted the residuals of this equation for each global and regional measure. We then performed a Levene's test (F-test) on these residuals and calculated variance ratios as Female SD / Male SD. The significance threshold was set to 0.05/306.

The AIs of all volumes were more variable in females compared to males in 3% (4/148) of regions and were more variable in males in 55% (82/148). The AIs of cortical surface areas were more variable in females compared to males in 5% (4/74) of regions and were more variable in males in 53% (39/74). The AIs of cortical mean thicknesses were more variable in females compared to males in 26% (19/74) of regions and were more variable in males in 15% (11/74; Supplemental Tables H1-4).

4. Reporting Raw AIs to Compare Results with Previous Studies.

**a) Methods**

We tested whether the raw AIs (Figure S4) were significantly different from 0 by performing a one-sample t-test (mu = 0) on each cortical AI in the UK Biobank for the Destrieux Segmentations and reported Cohen's D (Supplemental File S6; Supplemental Tables I1-6).

We additionally ran these analyses on the segmentations from the Desikan-Killiany-Trouville (DKT) atlas to compare results between atlases and to facilitate comparison with a previous study that analyzed segmentations from the Desikan-Killiany (DK) atlas (Kong et al., 2018). Kong and colleagues (2018) conducted the largest study examining sex and age effects on cortical brain asymmetries to date. They meta-analyzed the asymmetries of the DK mean thicknesses and surface areas in 17,141 individuals from the Enhancing NeuroImaging Genetics through Meta-Analysis (ENIGMA) Consortium and calculated the AI as AI = (L-R Measure) / ( (L+R Measure) / 2). We calculated the AIs of the UK Biobank Destrieux and DKT mean thicknesses and surface areas with their AI formula.

We then compared the subcortical AIs from our study to those from previous studies (Guadalupe et al., 2017; Sarica et al., 2018). Guadalupe and colleagues (2017) conducted the largest study examining sex and age effects on subcortical brain asymmetries to date. They meta-analyzed asymmetries across subcortical regions in 15 847 individuals in the ENIGMA Consortium and calculated the AI as AI = (L-R Measure) / (L+R Measure). Sarica and colleagues (2018) are the only ones to have examined asymmetries in hippocampal subfields and calculated the absolute AIs (AI = | L-R | / (L+R) ) of 100 healthy controls from the Alzheimer's Disease Neuroimaging Initiative (ADNI; Sarica et al., 2018)

Data from previous studies were taken from Supplemental Tables 2 for cortical regions (Kong et al., 2018), from Tables 2 for subcortical volumes (Guadalupe et al., 2017), and from Table 2 for the hippocampus and hippocampal subfields (Sarica et al., 2018).

**b) Results**

Destrieux volumes were asymmetric in all regions ($p < (0.05/74)$) except for the inferior part of the precentral sulcus. Destrieux surface areas were asymmetric in all regions except for the posterior-ventral part of the cingulate gyrus and mean thicknesses were asymmetric except for the frontal marginal gyrus and sulcus, the vertical ramus of the anterior segment of the lateral sulcus (or fissure), the occipital pole, and the anterior transverse collateral sulcus. There were leftward asymmetries in 45% (33/74) volumes, 47% (35/74) mean thicknesses, and 47% (35/74) surface areas, and a rightward asymmetry in 54% (40/74) volumes, 47% (35/74) mean thicknesses, L-R, and 49% (36/74) surface areas (Figure S5). The mean AI for volumes was 2.54e-4 (SD = 0.09), -9.67e-4 for mean thicknesses (SD = 0.01), and 1.34e-3 for surface areas (SD = 0.09).

In contrast, the surface areas and volumes of the DKT Segmentation Atlases were all asymmetric (p<(0.05/31)). The pars orbitalis and posterior cingulate gyri mean thicknesses were asymmetric according to the DK segmentations (Supplemental File S6; Figure S6).

The accumbens area, amygdala, caudate, and hippocampus had rightward asymmetries in the ENIGMA Consortium and UK Biobank, and the putamen, thalamus, and pallidum had leftwards asymmetries in the ENIGMA Consortium and leftward asymmetries or null asymmetries in the UK Biobank (Supplemental File S8).

Finally, the absolute hippocampal AIs (AI = | L-R | / (L+R) ) of 100 healthy controls from the ADNI (Sarica et al., 2018) were generally consistent with those from the UK Biobank, except for the parasubiculum which was symmetric in the UK Biobank and largely asymmetric in the ADNI (Supplemental File S9).

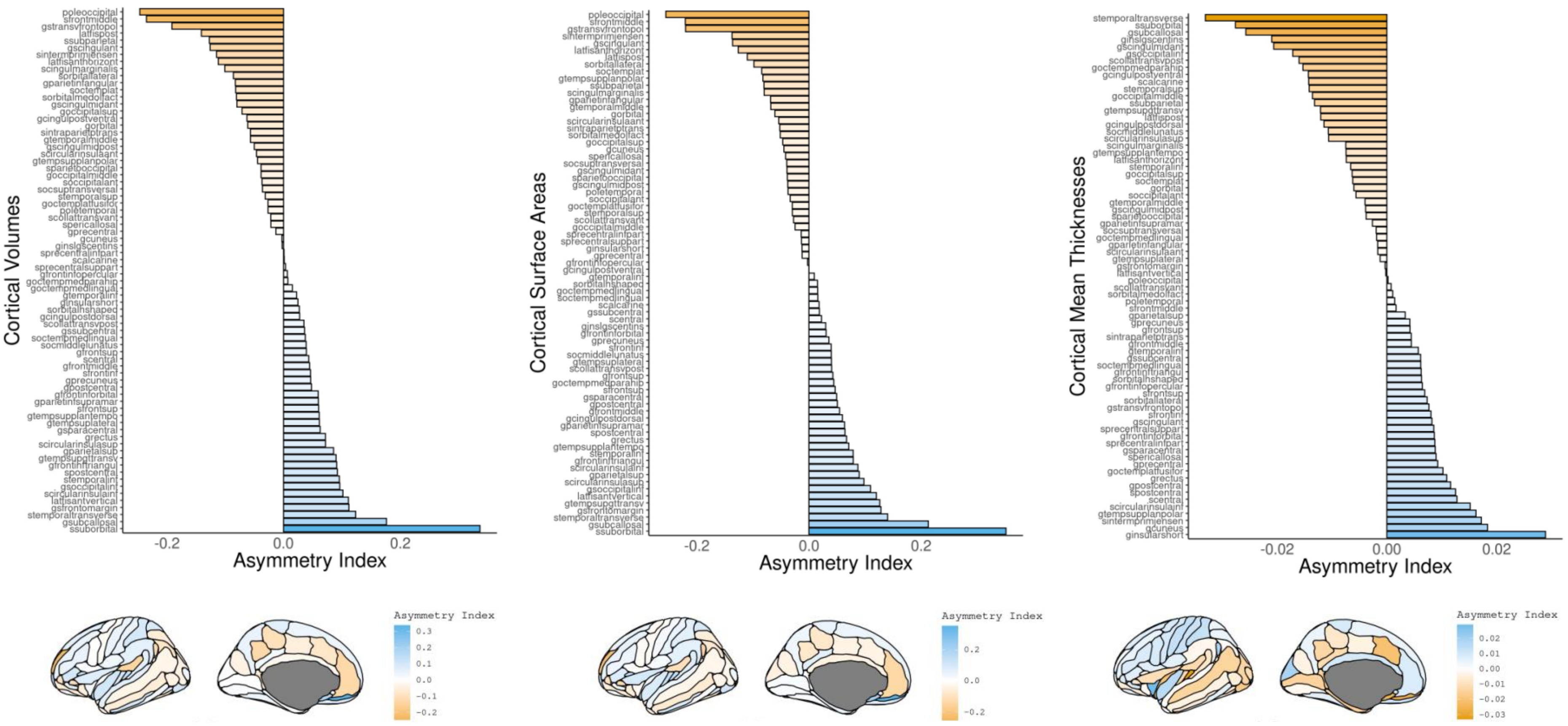


***Figure S5***. *Raw Asymmetry Index (AI) across Cortical Volumes, Surface Areas, and Mean Thicknesses*. Light Blue Regions: Left > Right Measure and Orange Regions: Right > Left Measure. AIs correspond to AI = (L-R) / ((L+R) / 2). Figures made with ggseg (Mowinckel & Vidal-Piñeiro, 2019).

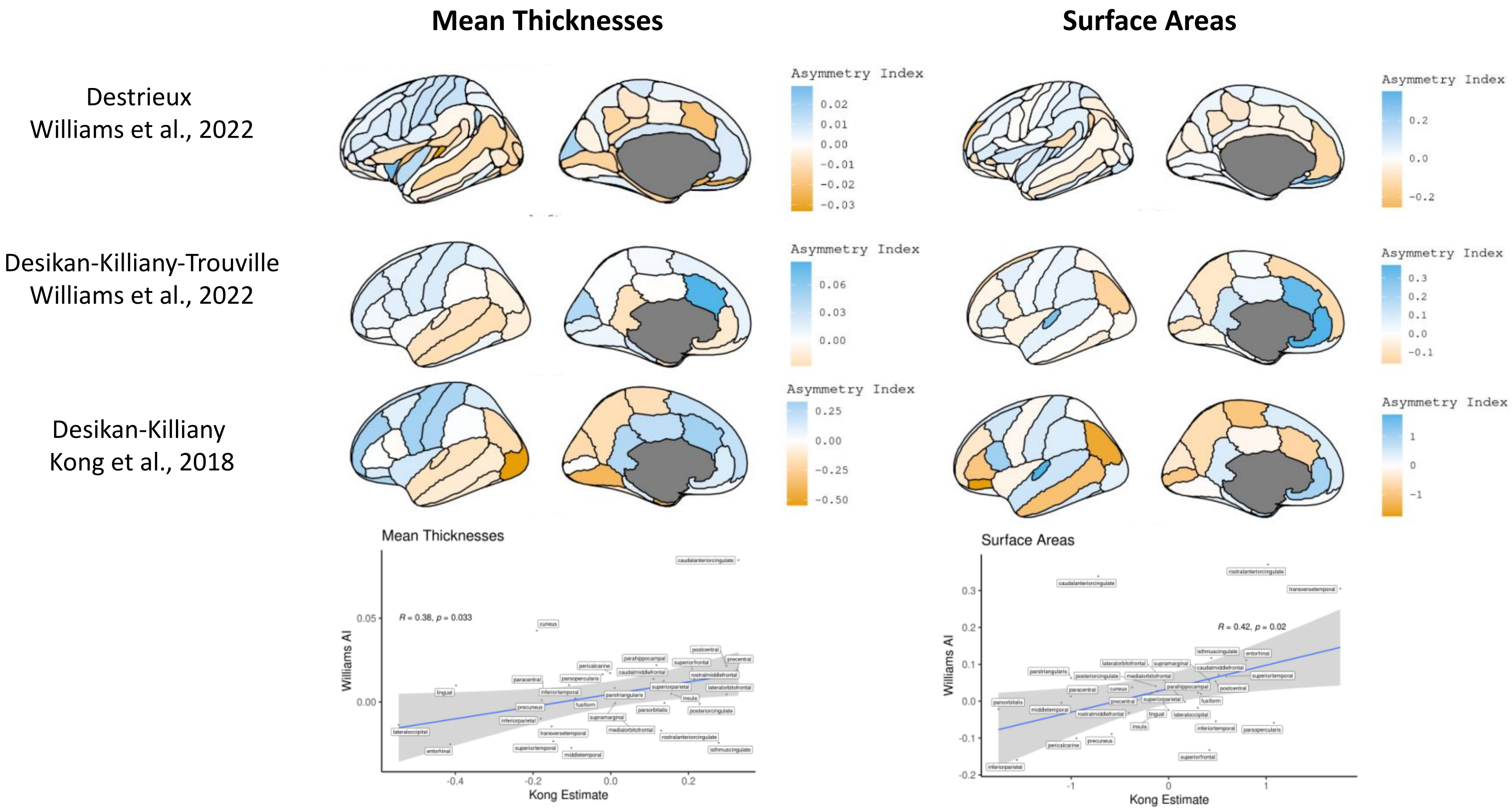


***Figure S6***. *Correlation between the magnitude of the asymmetry across Desikan-Killiany-Trouville segmentations from the UK Biobank (Williams) and the Desikan-Killiany segmentations ENIGMA cohort (Kong).* Light Blue Regions: Left > Right Measure and Orange Regions: Right > Left Measure. AIs from Williams et al., correspond to AI = (L-R) / ((L+R) / 2). Estimate values were taken from the meta-analysis of AIs ( (L-R) / ((L+R) / 2) ) from Kong et al., 2018

### c) Discussion - Cortical Regions

We observed a leftward asymmetry in frontal and parietal regions and a rightward asymmetry in temporal and occipital regions across volumes, mean thicknesses, and surface areas in the UK Biobank. More recent studies support the presence of this pattern across mean thicknesses (Kong et al., 2018; Plessen et al., 2014), although it was absent in surface areas in the latest large-scale meta-analysis on cerebral asymmetries (Kong et al., 2018). Differences in raw cortical AIs may be due to variations in image acquisition, image processing, and sampling (e.g., ENIGMA Consortium studies meta-analyzed data from individuals between 10 and 78 years old; Kong et al., 2018).

Several studies proposed that leftward structural asymmetries in the language network may mirror the leftward language dominance observed across functional studies (Bishop, 2013; Hagoort, 2005; Mazoyer et al., 2014). We indeed find leftward structural asymmetries in the language network, such as the well-established leftward asymmetry in volumes and surface areas in the planum temporale (Guadalupe et al., 2015; Kong et al., 2018; Maingault et al., 2016; Tzourio-Mazoyer et al., 2018). Consistent with Kong and colleagues' (2018) results, we additionally report a leftward asymmetry of the anterior transverse temporal gyrus of Heschl.

In line with a study on 215 healthy participants (Plessen et al., 2014), we also observe a leftward asymmetry in the mean thicknesses of the inferior frontal cortex (including Broca's area). However, a study on 250 adults reported a rightward asymmetry in these regions (Maingault et al., 2016) and an ENIGMA meta-analysis found that the pars opercularis and pars triangularis regions were symmetric and that the pars orbitalis had a leftward asymmetry (Kong et al., 2018). In our study, we saw the opposite pattern to that of the ENIGMA study: a leftward asymmetry in the pars opercularis and pars triangularis regions and symmetry in the pars orbitalis across the DKT and Destrieux segmentations. We also found a rightward asymmetry in the mean thickness of the supramarginal gyrus with the DKT atlas, whereas previous studies reported a leftward asymmetry (Plessen et al., 2014) or symmetry in this region (Koelkebeck et al., 2014; Kong et al., 2018). The Destrieux segmentations revealed a mix of rightward, leftward, and the absence of asymmetry in these regions, which could explain differences in regional AIs across studies.

Some researchers propose that either asymmetry in cortical thicknesses (Plessen et al., 2014) or surface areas (Kong et al., 2018) could anatomically correspond to functional language lateralization in these regions. Yet, we find rightward and leftward asymmetry in mean thicknesses and in surface areas of regions involved in language. While both asymmetries in surfaces and cortical thicknesses may contribute to language lateralization, language dominance observed across functional studies could also be independent of structural variations (Mazoyer et al., 2014).

#### d) Discussion - Subcortical Regions

The AIs of subcortical volumes were generally consistent with previous studies on brain asymmetries. In line with Guadalupe and colleagues' (2017) work on the ENIGMA Consortium, we found that the accumbens area, amygdala, caudate, and hippocampus had rightward asymmetries, whereas the putamen, thalamus, and pallidum had leftwards asymmetries. These findings also largely mirror those from a meta-analysis performed in a Japanese consortium, the COCORO (Cognitive Genetics Collaborative Research Organization; Healthy Control N = 1,680), although they reported that the pallidum and the caudate had a greater leftward asymmetry in controls (Okada et al., 2016).

### 5. Comparing Cortical Age and Sex Effects

#### a. Methods

Kong and colleagues (2018) and reported pooled sex and age effects from 17,141 individuals in the ENIGMA Consortium. They controlled for differences in brain size by adjusting for the Intracranial Volume (ICV) and calculated the AI as AI = (L-R Measure) / ( (L+R Measure) / 2). Their analyses included sex, linear age, quadratic age, ICV, and handedness (see their supplemental information here). Data from Kong and colleagues (2018) were taken from Supplemental Tables 3 and 4.

To compare their sex and age effects to those obtained with our models, we calculated the AIs of the DKT and Destrieux mean thicknesses and surface areas with their AI formula (Figure S5) and applied equation 12 to these AIs. Differences in the significance of sex and age estimates across studies could be explained by different raw AIs, segmentation algorithms, statistical models, and sample characteristics (e.g., age).

We tested whether differences in significant sex and age effects on brain asymmetries according to Kong and colleagues (2018) were due to brain measures that were included in our main analyses (i.e., L+R Measure and TCM AI). We ran equation S1 (equation 12 from the main text) on the Destrieux and DKT segmentations. We then ran equation S2 and S3 on the DKT segmentations. We compared the significant results from equation S1 with those from equation S2 to examine the effects of omitting L+R and L-R TCM. We compared the results of equation S3 to those reported by Kong and colleagues (2018) who controlled for ICV instead of TCM. TBV and ICV are highly correlated and are expected to yield similar results. Adjustment for multiple comparison was set at $p < (0.05/(306*5))$ for the UK Biobank.

**Equation S1**: AI = Intercept + ß$_1$*(L+R Measure)
+ ß$_2$*(TCM)
+ ß$_3$*(L-R TCM)
+ ß$_4$*(Sex) + ß$_5$*(age) + ß$_6$*(age$^2$)
+ ß$_7$*(sex) x (age) + ß$_8$*(sex) x (age$^2$)
+ ß$_9$*(handedness) + ß$_{10}$*(Scanner Site) + error

**Equation S2**: AI = Intercept + ß$_1$* (TCM)
+ ß$_2$*(Sex) + ß$_3$*(age) + ß$_4$*(age$^2$)
+ ß$_5$*(sex) x (age) + ß$_6$*(sex) x (age$^2$)
+ ß$_7$*(handedness) + ß$_8$*(Scanner Site) + error

**Equation S3**: AI = Intercept + $ß_1$* (TBV)

+ $ß_2$*(Sex) + $ß_3$*(age) + $ß_4$*($age^2$)

+ $ß_5$*(sex) x (age) + $ß_6$*(sex) x ($age^2$)

+ $ß_7$*(handedness) + $ß_8$*(Scanner Site) + error

#### b) Results - Cortical Sex Differences in AIs

Sex differences in surface area asymmetry were consistent across the ENIGMA Consortium (DK atlas) and the UK Biobank (Destrieux and DKT atlases), specifically across the inferior parietal, superior temporal, parahippocampal, and supramarginal surface areas. We found a sex difference in asymmetry in the superior frontal surface areas reported by Kong and colleagues (2018) in the Destrieux but not the DKT segmentations. In line with Kong and colleagues (2018), we found sex differences indicating greater female asymmetry in the caudal anterior cingulate and rostral anterior cingulate surface area asymmetries (Supplemental File S7), although they were not significant in our sample. In contrast to Kong and colleagues (2018), we did not find a sex difference in the lateral orbitofrontal surface areas in the DKT segmentations. This could be explained by the difference in the raw AI, as Kong and colleagues (2018) report a more leftward asymmetry and we find a greater rightward asymmetry in this region (Figure S7).

Kong and colleagues (2018) reported a sex difference in the asymmetry of the entorhinal and the parahippocampal gyrus mean thicknesses. We did not find a sex difference in these regions in our replication of their analyses with the DKT atlas, although the sex difference in the asymmetry of the entorhinal cortex mean thickness became significant when adjusting for TCM, L-R TCM, and L+R.

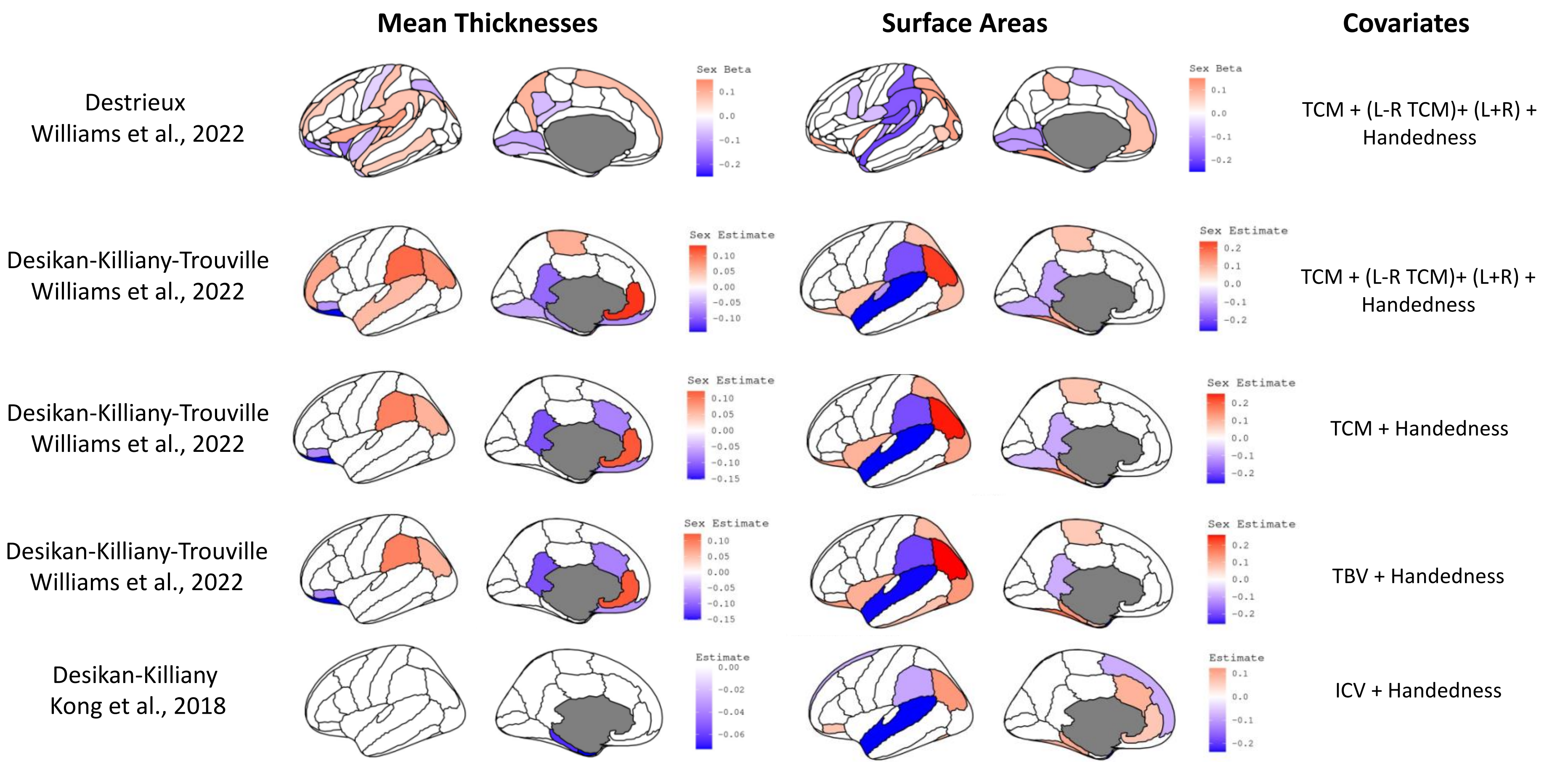


***Figure S7****. Significant Sex Effects in Cortical Asymmetries across Destrieux and Desikan-Killiany-Trouville Segmentations from the UK Biobank (Williams) and the Desikan-Killiany segmentations from the ENIGMA consortium (Kong) with different covariates.* All Williams and colleagues' (2022) models also include scanner site and age by sex and age2 by sex interactions. Kong and colleagues (2018) meta-analyzed sex, linear age, and quadratic age effects. TCM: Total Mean Cortical Thickness for mean thicknesses, Total Surface Area for surface areas, TBV: Total Brain Volume, ICV: Intracranial Volume.

#### c) Results - Comparing Cortical Age Differences in AIs

In line with Kong and colleagues (2018), we found a greater leftward asymmetry with increased age in the mean thickness of the superior temporal gyrus. However, instead of reporting an increase in the rightward asymmetry with age in the entorhinal cortex, we found that the rightward asymmetry decreased with age (Figure S8).

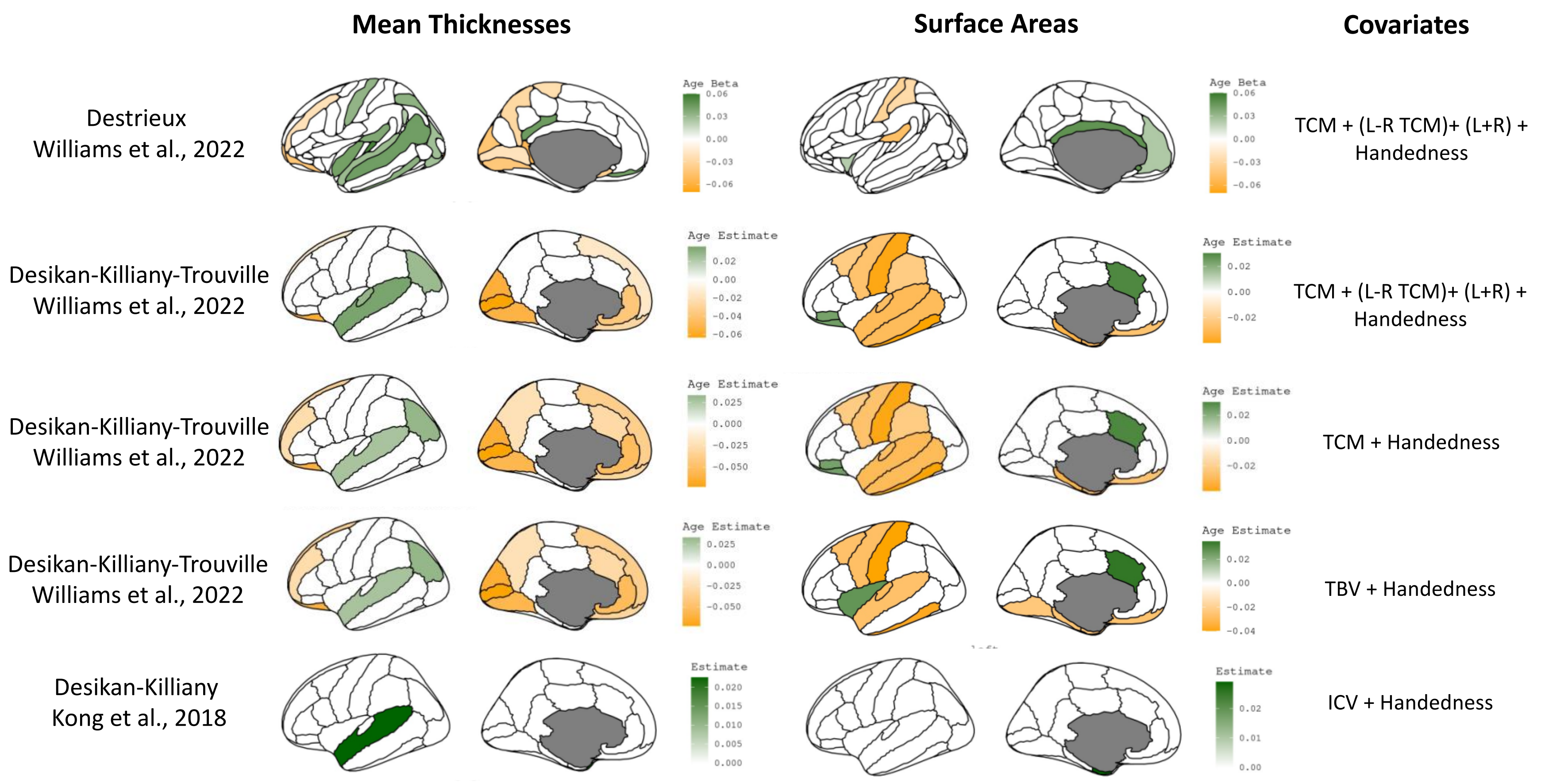


***Figure S8**. Significant Age Effects in Cortical Asymmetries across Destrieux and Desikan-Killiany-Trouville Segmentations from the UK Biobank (Williams) and the Desikan-Killiany segmentations from the ENIGMA consortium (Kong) with different covariates.* All Williams and colleagues (2022) models also include scanner site and age by sex and age2 by sex interactions. Kong and colleagues (2018) meta-analyzed sex, linear age, and quadratic age effects. TCM: Total Mean Cortical Thickness for mean thicknesses, Total Surface Area for surface areas, TBV: Total Brain Volume, ICV: Intracranial Volume.

### 6) Comparing Subcortical AIs and Sex and Age Effects

#### a) Methods

Guadalupe and colleagues (2017) tested for sex differences on the residualized AI, where linear age, ICV, and handedness were removed, and for the meta-analyses of age effects, they used the age coefficients from the ANCOVA with ICV and sex as covariates.

#### b) Results

Differences in AI may explain differences in the age and sex effects reported between our study and that of Guadalupe and colleagues (2017; Figure S9).

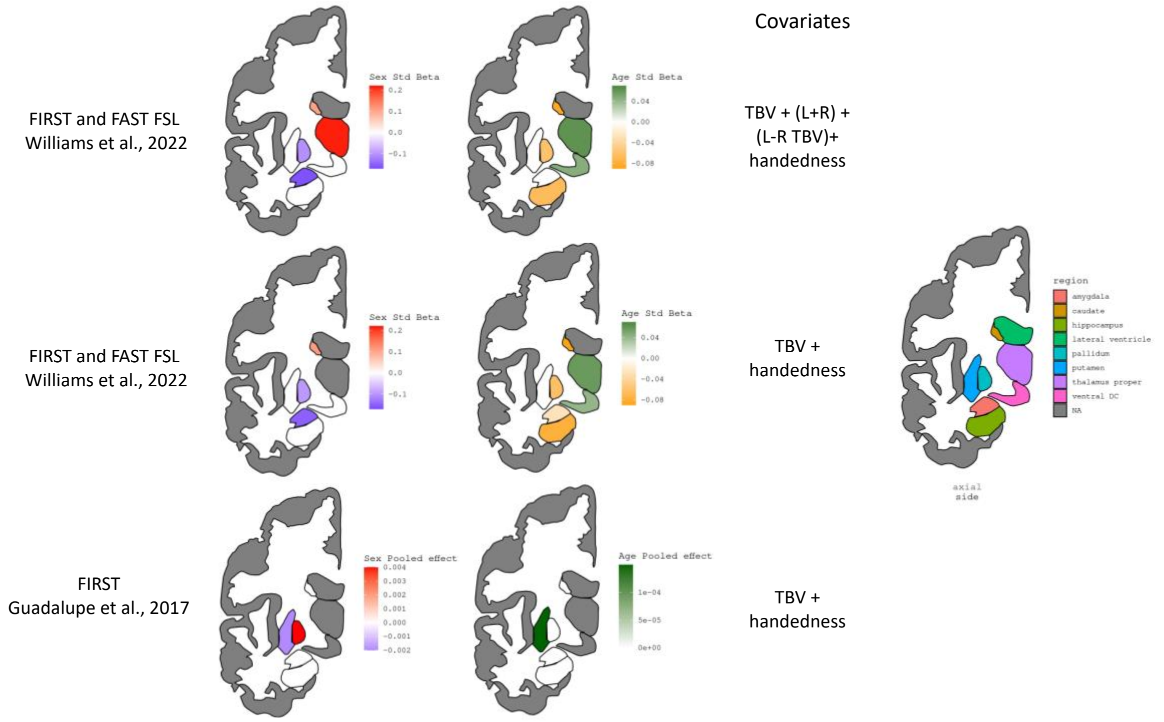


***Figure S9****. Significant Subcortical Age and Sex Effects between the Asymmetry Index (AI) and the AI Sex and Age Effect in the UK Biobank (Williams) and the ENIGMA consortium (Guadalupe).* Sex and age effects correspond to standardized betas for the UK Biobank and pooled effects for the ENIGMA consortium. All Williams and colleagues' present models also include scanner site and age by sex and age2 by sex interactions. Kong and colleagues (2018) effects include ICV, handedness, age, and sex. TBV: Total Brain Volume.

### 6 - R packages

The following R packages were used in the study:

- data.table (Dowle et al., 2020)
- ggplot2 (Wickham, 2016)
- tidyr (Wickham & RStudio, 2020)
- broom (Robinson et al., 2020)
- sjPLot (Lüdecke et al., 2020)
- car (Fox et al., 2020)
- ggpubr (Kassambara, 2020)
- plyr (Wickham, 2020)
- dplyr (Wickham, François, et al., 2020)
- Tables (Murdoch, 2020)
- Scales (Wickham, Seidel, et al., 2020)
- Corrplot (Wei et al., 2017)
- Ggrepel (Slowikowski et al., 2020)
- Ggeseg (Mowinckel & Vidal-Piñeiro, 2019)
