## Supplementary material for "Comparing Brain Asymmetries Independently of Brain Size": File_S1_Q2_Non_Linear_Global_Measures.html


### Non\_Linear\_Global\_Measures

### Table of All Regions with a reduction in RSE > 1%

```
##        effect                 region percent_change percent_change_sig
##  1:    poly 3           Cortical GMV       1.172822                  1
##  2:        bs           Cortical GMV       1.172822                  1
##  3: smooth tp           Cortical GMV       1.354692                  1
##  4:    poly 3            Cerebral WM       1.350478                  1
##  5:        bs            Cerebral WM       1.350478                  1
##  6: smooth tp            Cerebral WM       1.816211                  1
##  7:    poly 2          volume of lsg       1.510965                  1
##  8:    poly 3          volume of lsg       3.123744                  1
##  9:        bs          volume of lsg       3.123744                  1
## 10:      ns 2          volume of lsg       1.625199                  1
## 11:      ns 3          volume of lsg       3.674838                  1
## 12: smooth tp          volume of lsg       3.815887                  1
## 13:    poly 3  volume of ssuborbital       1.427064                  1
## 14:        bs  volume of ssuborbital       1.427064                  1
## 15:      ns 3  volume of ssuborbital       1.443774                  1
## 16: smooth tp  volume of ssuborbital       1.252216                  1
## 17:    poly 2 area of ginslgscentins       1.248228                  1
## 18:    poly 3 area of ginslgscentins       1.386710                  1
## 19:        bs area of ginslgscentins       1.386710                  1
## 20:      ns 2 area of ginslgscentins       1.318630                  1
## 21:      ns 3 area of ginslgscentins       1.356951                  1
## 22: smooth tp area of ginslgscentins       1.196950                  1
## 23:    poly 2    area of ssuborbital       2.168808                  1
## 24:    poly 3    area of ssuborbital       2.297415                  1
## 25:        bs    area of ssuborbital       2.297415                  1
## 26:      ns 2    area of ssuborbital       2.222919                  1
## 27:      ns 3    area of ssuborbital       2.306610                  1
## 28: smooth tp    area of ssuborbital       2.016991                  1
##        effect                 region percent_change percent_change_sig
```

### Plots of Global Measures

#### 1. Cortical Grey Matter Volume (GMV)

#### 2. Cerebral White Matter Volume (WMV)

### II - Exploratory Models

#### Residual Standard Error (RSE) % Change by Model and Region

###### Models

We conducted exploratory analyses to identify the cerebral predictors responsible for the reduction in RSE in these regions. We ran the three models for each region. Each model consisted of the cerebral AI as a function of either the Left + Right Measure, the Total Cerebral Measure, or the Total Cerebral Measure AI.

poly 3: polynomial to the third degree

bs: B-spline basis matrix for a polynomial spline (df = 3)

tp: thin plate smooth spline

```
##   Region and Model Predictor      poly 3          bs           tp
## 1        Cortical_GMV_IV_SUM 0.004534119 0.004534119 1.129304e-07
## 2        Cortical_GMV_TBV_AI 1.110442507 1.110442507 1.523599e+00
## 3           Cortical_GMV_TBV 0.006055990 0.006055990 3.639884e-03
## 4        Cerebral_WMV_IV_SUM 0.008294602 0.008294602 9.831114e-03
## 5        Cerebral_WMV_TBV_AI 1.353114443 1.353114443 1.827780e+00
## 6           Cerebral_WMV_TBV 0.012199395 0.012199395 1.440609e-02
```
