## Supplementary material for "Comparing Brain Asymmetries Independently of Brain Size": File_S2_Q2_Non_Linear_Surface_Areas.html

Q2\_Non\_Linear\_SA\_Plots


### Q2\_Non\_Linear\_SA\_Plots

### Table of Regions with a reduction in RSE > 1%

```
##        effect         region percent_change percent_change_sig
##  1:    poly 2 ginslgscentins       1.248228                  1
##  2:    poly 3 ginslgscentins       1.386710                  1
##  3:   poly bs ginslgscentins       1.386710                  1
##  4: poly ns 2 ginslgscentins       1.318630                  1
##  5: poly ns 3 ginslgscentins       1.356951                  1
##  6: smooth tp ginslgscentins       1.196950                  1
##  7:    poly 2    ssuborbital       2.168808                  1
##  8:    poly 3    ssuborbital       2.297415                  1
##  9:   poly bs    ssuborbital       2.297415                  1
## 10: poly ns 2    ssuborbital       2.222919                  1
## 11: poly ns 3    ssuborbital       2.306610                  1
## 12: smooth tp    ssuborbital       2.016991                  1
```

### Plots

#### 1. ginslgscentins - Long insular gyrus and central sulcus of the insula

#### 2. ssuborbital - suborbital sulcus

### II - Exploratory Models

#### Residual Standard Error (RSE) % Change by Model and Region

###### Models

We conducted exploratory analyses to identify the cerebral predictors responsible for the reduction in RSE in these regions. We ran the three models for each region. Each model consisted of the cerebral AI as a function of either the Left + Right Measure, the Total Cerebral Measure, or the Total Cerebral Measure AI.

poly 2: polynomial to the second degree

poly 3: polynomial to the third degree

bs: B-spline basis matrix for a polynomial spline (df = 3)

ns 2: natural spline (df = 2)

ns 3: natural spline (df = 3)

tp: thin plate smooth spline

```
##           Region and Model Predictor        poly 2        poly 3            bs
## 1              ginslgscentins_IV_SUM  1.1359401816  1.2384411659  1.2384411659
## 2 ginslgscentins_Cortical_TSA_sum_AI  0.0050262844  0.0045391514  0.0045391514
## 3    ginslgscentins_Cortical_TSA_sum -0.0012426496 -0.0010119673 -0.0010119673
## 4                 ssuborbital_IV_SUM  2.0201044602  2.1341801891  2.1341801891
## 5    ssuborbital_Cortical_TSA_sum_AI  0.0009038613  0.0001090024  0.0001090024
## 6       ssuborbital_Cortical_TSA_sum  0.0079923058  0.0070689708  0.0070689708
##            ns 2          ns 3           tp
## 1  1.1957728062  1.2140920750 1.2948054576
## 2  0.0068475462  0.0086718548 0.0174479913
## 3 -0.0012596956  0.0021996903 0.0070418689
## 4  2.0707238340  2.1436354889 2.1706856680
## 5  0.0007924523 -0.0004817951 0.0001156642
## 6  0.0082574533  0.0085059226 0.0209535835
```
