## Supplementary material for "Comparing Brain Asymmetries Independently of Brain Size": File_S3_Q2_Non_Linear_Volumes.html


### Non\_linear\_volumes

### Table of Regional Volumes with a reduction in RSE > 1%

```
##        effect      region percent_change percent_change_sig
##  1:    poly 2         lsg       1.510965                  1
##  2:    poly 3         lsg       3.123744                  1
##  3:   poly bs         lsg       3.123744                  1
##  4: poly ns 2         lsg       1.625199                  1
##  5: poly ns 3         lsg       3.674838                  1
##  6: smooth tp         lsg       3.815887                  1
##  7:    poly 3 ssuborbital       1.427064                  1
##  8:   poly bs ssuborbital       1.427064                  1
##  9: poly ns 3 ssuborbital       1.443774                  1
## 10: smooth tp ssuborbital       1.252216                  1
```

### Plots

#### 1. lsg - limitans suprageniculate thalamic nuclei

#### 2. ssuborbital - suborbital sulcus

### II - Exploratory Models

#### Residual Standard Error (RSE) % Change by Model and Region

###### Models

We conducted exploratory analyses to identify the cerebral predictors responsible for the reduction in RSE in these regions. We ran the three models for each region. Each model consisted of the cerebral AI as a function of either the Left + Right Measure, the Total Cerebral Measure, or the Total Cerebral Measure AI.

poly 2: polynomial to the second degree

poly 3: polynomial to the third degree

bs: B-spline basis matrix for a polynomial spline (df = 3)

ns 2: natural spline (df = 2)

ns 3: natural spline (df = 3)

tp: thin plate smooth spline

```
##   Region and Model Predictor      poly 2      poly 3          bs        ns 2
## 1                 lsg_IV_SUM 1.524806435 3.155573209 3.155573209 1.639954037
## 2                 lsg_TBV_AI 0.032852962 0.031575092 0.031575092 0.033583795
## 3                    lsg_TBV 0.001334886 0.005862129 0.005862129 0.001145095
##         ns 3         tp
## 1 3.69246511 3.84958045
## 2 0.03599020 0.05628597
## 3 0.01229991 0.01278611
```

```
##   Region and Model Predictor      poly 3          bs          ns 3
## 1         ssuborbital_IV_SUM 1.488321453 1.488321453  1.5057689522
## 2         ssuborbital_TBV_AI 0.001915696 0.001915696 -0.0006829684
## 3            ssuborbital_TBV 0.012846759 0.012846759  0.0141783243
##              tp
## 1  1.525119e+00
## 2 -1.078345e-10
## 3  2.070227e-02
```
