## Supplementary material for "Comparing Brain Asymmetries Independently of Brain Size": File_S4_Q3_on_Non_Linear_Models.html

Q3\_Non\_Linear\_Measures


### Q3\_Non\_Linear\_Measures

# I - SA

```
DT1<-fread("/scratch2/ukbio/biobank/Asymmetry/Asymmetry_2/data/AI_SA_a2009s.csv")
DT <- as.data.frame(DT1)

scanner_data <-fread("/scratch2/ukbio/biobank/Allometry/data/site_scanner_df.csv") 
scanner_data <- as.data.frame(scanner_data)
Merged_DF <- left_join(DT, scanner_data, by = "eid")
Merged_DF$Scanner_Site <- as.factor(Merged_DF$uk_biobank_assessment_centre_f54_2_0)
table(Merged_DF$Scanner_Site)
```

```
## 
## 11025 11026 11027 
## 23766  5042  9901
```

```
DT <- Merged_DF

scanner_data <-fread("/scratch2/ukbio/biobank/Asymmetry/Asymmetry_2/data/Global_Measures_AI.csv")
scanner_data <- as.data.frame(scanner_data)
Merged_DF <- left_join(DT, scanner_data)
```

```
## Joining, by = c("eid", "TBV_log", "TBV", "Age_MRI_1", "sex")
```

```
DT1 <- Merged_DF

# area of ginslgscentins L +R 
Non_Linear_Model <- gam(scale(area_of_ginslgscentins_AI_old) ~ bs(scale(area_of_ginslgscentins_sum))+ 
                   scale(Cortical_TSA_AI_regional) + scale(Cortical_TSA_sum) +  
                   sex +
                   Scanner_Site, data = DT1)
summary(Non_Linear_Model)
```

```
## 
## Family: gaussian 
## Link function: identity 
## 
## Formula:
## scale(area_of_ginslgscentins_AI_old) ~ bs(scale(area_of_ginslgscentins_sum)) + 
##     scale(Cortical_TSA_AI_regional) + scale(Cortical_TSA_sum) + 
##     sex + Scanner_Site
## 
## Parametric coefficients:
##                                         Estimate Std. Error t value Pr(>|t|)
## (Intercept)                            -0.994702   0.074348 -13.379  < 2e-16
## bs(scale(area_of_ginslgscentins_sum))1  2.842984   0.187174  15.189  < 2e-16
## bs(scale(area_of_ginslgscentins_sum))2 -0.425291   0.109423  -3.887 0.000102
## bs(scale(area_of_ginslgscentins_sum))3 -1.005178   0.222372  -4.520 6.19e-06
## scale(Cortical_TSA_AI_regional)         0.052399   0.005014  10.450  < 2e-16
## scale(Cortical_TSA_sum)                 0.076691   0.007140  10.742  < 2e-16
## sex                                     0.142522   0.011990  11.887  < 2e-16
## Scanner_Site11026                      -0.008263   0.015098  -0.547 0.584191
## Scanner_Site11027                      -0.002181   0.011678  -0.187 0.851867
##                                           
## (Intercept)                            ***
## bs(scale(area_of_ginslgscentins_sum))1 ***
## bs(scale(area_of_ginslgscentins_sum))2 ***
## bs(scale(area_of_ginslgscentins_sum))3 ***
## scale(Cortical_TSA_AI_regional)        ***
## scale(Cortical_TSA_sum)                ***
## sex                                    ***
## Scanner_Site11026                         
## Scanner_Site11027                         
## ---
## Signif. codes:  0 '***' 0.001 '**' 0.01 '*' 0.05 '.' 0.1 ' ' 1
## 
## 
## R-sq.(adj) =  0.0551   Deviance explained = 5.53%
## GCV = 0.94516  Scale est. = 0.94494   n = 38709
```

```
Beta1 <- 0.142522 
SE1 <- 0.011990

Linear_Model <- lm(scale(area_of_ginslgscentins_AI_old) ~ (scale(area_of_ginslgscentins_sum))+ 
                   scale(Cortical_TSA_AI_regional) + scale(Cortical_TSA_sum) +  
                   sex +
                   Scanner_Site, data = DT1)
summary(Linear_Model)
```

```
## 
## Call:
## lm(formula = scale(area_of_ginslgscentins_AI_old) ~ (scale(area_of_ginslgscentins_sum)) + 
##     scale(Cortical_TSA_AI_regional) + scale(Cortical_TSA_sum) + 
##     sex + Scanner_Site, data = DT1)
## 
## Residuals:
##     Min      1Q  Median      3Q     Max 
## -4.4252 -0.6232  0.0612  0.6886  3.8627 
## 
## Coefficients:
##                                    Estimate Std. Error t value Pr(>|t|)    
## (Intercept)                       -0.002015   0.006403  -0.315    0.753    
## scale(area_of_ginslgscentins_sum) -0.186487   0.006296 -29.622   <2e-16 ***
## scale(Cortical_TSA_AI_regional)    0.053187   0.005074  10.481   <2e-16 ***
## scale(Cortical_TSA_sum)            0.098899   0.007187  13.761   <2e-16 ***
## sex                                0.142177   0.012130  11.721   <2e-16 ***
## Scanner_Site11026                 -0.009224   0.015278  -0.604    0.546    
## Scanner_Site11027                 -0.003686   0.011818  -0.312    0.755    
## ---
## Signif. codes:  0 '***' 0.001 '**' 0.01 '*' 0.05 '.' 0.1 ' ' 1
## 
## Residual standard error: 0.9837 on 38702 degrees of freedom
## Multiple R-squared:  0.03241,    Adjusted R-squared:  0.03226 
## F-statistic: 216.1 on 6 and 38702 DF,  p-value: < 2.2e-16
```

```
Beta2 <- 0.142177 
SE2 <- 0.012130

Z = ( Beta1 - Beta2) / sqrt((SE1^2)+ (SE2^2))
Z
```

```
## [1] 0.02022784
```

```
# area_of_ssuborbital L +R 
Non_Linear_Model <- gam(scale(area_of_ssuborbital_AI_old) ~ s(scale(area_of_ssuborbital_sum))+ 
                   scale(Cortical_TSA_AI_regional) + scale(Cortical_TSA_sum) +  
                   sex +Scanner_Site, data = DT1)
summary(Non_Linear_Model)
```

```
## 
## Family: gaussian 
## Link function: identity 
## 
## Formula:
## scale(area_of_ssuborbital_AI_old) ~ s(scale(area_of_ssuborbital_sum)) + 
##     scale(Cortical_TSA_AI_regional) + scale(Cortical_TSA_sum) + 
##     sex + Scanner_Site
## 
## Parametric coefficients:
##                                  Estimate Std. Error t value Pr(>|t|)    
## (Intercept)                     -0.008153   0.006249  -1.305  0.19203    
## scale(Cortical_TSA_AI_regional)  0.024069   0.004951   4.861 1.17e-06 ***
## scale(Cortical_TSA_sum)          0.082932   0.006629  12.511  < 2e-16 ***
## sex                              0.032826   0.011846   2.771  0.00559 ** 
## Scanner_Site11026                0.014068   0.014906   0.944  0.34528    
## Scanner_Site11027                0.020957   0.011534   1.817  0.06922 .  
## ---
## Signif. codes:  0 '***' 0.001 '**' 0.01 '*' 0.05 '.' 0.1 ' ' 1
## 
## Approximate significance of smooth terms:
##                                     edf Ref.df     F p-value    
## s(scale(area_of_ssuborbital_sum)) 5.833  6.998 446.4  <2e-16 ***
## ---
## Signif. codes:  0 '***' 0.001 '**' 0.01 '*' 0.05 '.' 0.1 ' ' 1
## 
## R-sq.(adj) =  0.0783   Deviance explained = 7.85%
## GCV = 0.92202  Scale est. = 0.92174   n = 38709
```

```
Linear_Model <- lm(scale(area_of_ssuborbital_AI_old) ~ scale(area_of_ssuborbital_sum)+ 
                     scale(Cortical_TSA_AI_regional) + scale(Cortical_TSA_sum) +  
                     sex +Scanner_Site, data = DT1)
summary(Linear_Model)
```

```
## 
## Call:
## lm(formula = scale(area_of_ssuborbital_AI_old) ~ scale(area_of_ssuborbital_sum) + 
##     scale(Cortical_TSA_AI_regional) + scale(Cortical_TSA_sum) + 
##     sex + Scanner_Site, data = DT1)
## 
## Residuals:
##     Min      1Q  Median      3Q     Max 
## -3.6582 -0.6823 -0.0603  0.6341  3.8081 
## 
## Coefficients:
##                                  Estimate Std. Error t value Pr(>|t|)    
## (Intercept)                     -0.007562   0.006391  -1.183  0.23676    
## scale(area_of_ssuborbital_sum)  -0.212712   0.005890 -36.112  < 2e-16 ***
## scale(Cortical_TSA_AI_regional)  0.024311   0.005064   4.801 1.58e-06 ***
## scale(Cortical_TSA_sum)          0.074775   0.006776  11.035  < 2e-16 ***
## sex                              0.031428   0.012109   2.595  0.00945 ** 
## Scanner_Site11026                0.011896   0.015243   0.780  0.43516    
## Scanner_Site11027                0.019910   0.011795   1.688  0.09141 .  
## ---
## Signif. codes:  0 '***' 0.001 '**' 0.01 '*' 0.05 '.' 0.1 ' ' 1
## 
## Residual standard error: 0.9819 on 38702 degrees of freedom
## Multiple R-squared:  0.0361, Adjusted R-squared:  0.03595 
## F-statistic: 241.6 on 6 and 38702 DF,  p-value: < 2.2e-16
```

```
Beta1 <- 0.032826 
SE1 <- 0.011846
Beta2 <- 0.031428 
SE2 <- 0.015243

Z = ( Beta1 - Beta2) / sqrt((SE1^2)+ (SE2^2))
Z
```

```
## [1] 0.07241708
```

### II - Volumes

```
## Load Data
DT1<-fread("/scratch2/ukbio/biobank/Asymmetry/Asymmetry_2/data/AI_ASEG.csv")
DT <-fread("/scratch2/ukbio/biobank/Asymmetry/Asymmetry_2/data/AI_Vol_FAST.csv")
DT1 <- left_join(DT1, DT, by = "eid")
DT <-fread("/scratch2/ukbio/biobank/Asymmetry/Asymmetry_2/data/AI_Vol_Subseg.csv")
DT1 <- left_join(DT1, DT, by = "eid")
DT <-fread("/scratch2/ukbio/biobank/Asymmetry/Asymmetry_2/data/AI_Vol_a2009s.csv")
DT1 <- left_join(DT1, DT, by = "eid")
DT <- as.data.frame(DT1)

## Add Scanner 
scanner_data <-fread("/scratch2/ukbio/biobank/Allometry/data/site_scanner_df.csv") 
scanner_data <- as.data.frame(scanner_data)
Merged_DF <- left_join(DT, scanner_data, by = "eid")
Merged_DF$Scanner_Site <- as.factor(Merged_DF$uk_biobank_assessment_centre_f54_2_0)
table(Merged_DF$Scanner_Site)
```

```
## 
## 11025 11026 11027 
## 24971  5070  9987
```

```
DT <- Merged_DF

## Add Left-Right TBV  
scanner_data <-fread("/scratch2/ukbio/biobank/Asymmetry/Asymmetry_2/data/Global_Measures_AI.csv")
scanner_data <- as.data.frame(scanner_data)
Merged_DF <- left_join(DT, scanner_data)
```

```
## Joining, by = "eid"
```

```
DT1 <- Merged_DF

# lsg L +R 
Non_Linear_Model <- gam(scale(volume_of_lsg_AI_old) ~ s(scale(volume_of_lsg_sum)) + scale(TBV_AI_regional) + 
                   scale(TBV) + 
                   sex +
                   Scanner_Site, data = DT1)
summary(Non_Linear_Model)
```

```
## 
## Family: gaussian 
## Link function: identity 
## 
## Formula:
## scale(volume_of_lsg_AI_old) ~ s(scale(volume_of_lsg_sum)) + scale(TBV_AI_regional) + 
##     scale(TBV) + sex + Scanner_Site
## 
## Parametric coefficients:
##                         Estimate Std. Error t value Pr(>|t|)    
## (Intercept)            -0.012156   0.006229  -1.951  0.05102 .  
## scale(TBV_AI_regional)  0.033681   0.004887   6.892 5.58e-12 ***
## scale(TBV)              0.057314   0.005963   9.612  < 2e-16 ***
## sex                     0.143720   0.011908  12.069  < 2e-16 ***
## Scanner_Site11026      -0.041569   0.014848  -2.800  0.00512 ** 
## Scanner_Site11027       0.050153   0.011485   4.367 1.26e-05 ***
## ---
## Signif. codes:  0 '***' 0.001 '**' 0.01 '*' 0.05 '.' 0.1 ' ' 1
## 
## Approximate significance of smooth terms:
##                               edf Ref.df     F p-value    
## s(scale(volume_of_lsg_sum)) 8.551  8.945 360.3  <2e-16 ***
## ---
## Signif. codes:  0 '***' 0.001 '**' 0.01 '*' 0.05 '.' 0.1 ' ' 1
## 
## R-sq.(adj) =  0.0845   Deviance explained = 8.48%
## GCV = 0.91585  Scale est. = 0.91551   n = 38709
```

```
Linear_Model <- lm(scale(volume_of_lsg_AI_old) ~ (scale(volume_of_lsg_sum)) + scale(TBV_AI_regional) + 
                          scale(TBV) + 
                          sex +
                          Scanner_Site, data = DT1)

summary(Linear_Model)
```

```
## 
## Call:
## lm(formula = scale(volume_of_lsg_AI_old) ~ (scale(volume_of_lsg_sum)) + 
##     scale(TBV_AI_regional) + scale(TBV) + sex + Scanner_Site, 
##     data = DT1)
## 
## Residuals:
##     Min      1Q  Median      3Q     Max 
## -4.9962 -0.6582 -0.0273  0.6649  3.6932 
## 
## Coefficients:
##                           Estimate Std. Error t value Pr(>|t|)    
## (Intercept)              -0.011543   0.006474  -1.783 0.074581 .  
## scale(volume_of_lsg_sum) -0.059222   0.005427 -10.913  < 2e-16 ***
## scale(TBV_AI_regional)    0.035567   0.005078   7.004 2.53e-12 ***
## scale(TBV)                0.079858   0.006177  12.928  < 2e-16 ***
## sex                       0.159935   0.012363  12.936  < 2e-16 ***
## Scanner_Site11026        -0.041768   0.015430  -2.707 0.006795 ** 
## Scanner_Site11027         0.045174   0.011934   3.785 0.000154 ***
## ---
## Signif. codes:  0 '***' 0.001 '**' 0.01 '*' 0.05 '.' 0.1 ' ' 1
## 
## Residual standard error: 0.9944 on 38702 degrees of freedom
##   (1319 observations deleted due to missingness)
## Multiple R-squared:  0.01141,    Adjusted R-squared:  0.01125 
## F-statistic: 74.43 on 6 and 38702 DF,  p-value: < 2.2e-16
```

```
Beta1 <- 0.143720 
SE1 <- 0.011908
Beta2 <- 0.159935 
SE2 <- 0.012363

Z = ( Beta1 - Beta2) / sqrt((SE1^2)+ (SE2^2))
Z
```

```
## [1] -0.9446436
```

```
# ssuborbital L +R 
Non_Linear_Model <- gam(scale(volume_of_ssuborbital_AI_old) ~ ns(scale(volume_of_ssuborbital_sum))+ scale(TBV_AI_regional) + 
                   scale(TBV) + 
                   sex +
                   Scanner_Site, data = DT1)
summary(Non_Linear_Model)
```

```
## 
## Family: gaussian 
## Link function: identity 
## 
## Formula:
## scale(volume_of_ssuborbital_AI_old) ~ ns(scale(volume_of_ssuborbital_sum)) + 
##     scale(TBV_AI_regional) + scale(TBV) + sex + Scanner_Site
## 
## Parametric coefficients:
##                                       Estimate Std. Error t value Pr(>|t|)    
## (Intercept)                          -0.458708   0.016761 -27.368  < 2e-16 ***
## ns(scale(volume_of_ssuborbital_sum))  1.906672   0.064981  29.342  < 2e-16 ***
## scale(TBV_AI_regional)                0.035140   0.005052   6.955 3.58e-12 ***
## scale(TBV)                           -0.060495   0.006274  -9.643  < 2e-16 ***
## sex                                   0.069384   0.012121   5.724 1.05e-08 ***
## Scanner_Site11026                    -0.005086   0.015356  -0.331   0.7405    
## Scanner_Site11027                     0.025424   0.011880   2.140   0.0323 *  
## ---
## Signif. codes:  0 '***' 0.001 '**' 0.01 '*' 0.05 '.' 0.1 ' ' 1
## 
## 
## R-sq.(adj) =  0.0243   Deviance explained = 2.45%
## GCV = 0.97584  Scale est. = 0.97566   n = 38564
```

```
Linear_Model <- lm(scale(volume_of_ssuborbital_AI_old) ~ (scale(volume_of_ssuborbital_sum))+ scale(TBV_AI_regional) + 
                          scale(TBV) + 
                          sex +
                          Scanner_Site, data = DT1)
summary(Linear_Model)
```

```
## 
## Call:
## lm(formula = scale(volume_of_ssuborbital_AI_old) ~ (scale(volume_of_ssuborbital_sum)) + 
##     scale(TBV_AI_regional) + scale(TBV) + sex + Scanner_Site, 
##     data = DT1)
## 
## Residuals:
##     Min      1Q  Median      3Q     Max 
## -4.0157 -0.6548 -0.0064  0.6389  3.2993 
## 
## Coefficients:
##                                   Estimate Std. Error t value Pr(>|t|)    
## (Intercept)                      -0.007066   0.006443  -1.097   0.2728    
## scale(volume_of_ssuborbital_sum)  0.157631   0.005372  29.342  < 2e-16 ***
## scale(TBV_AI_regional)            0.035140   0.005052   6.955 3.58e-12 ***
## scale(TBV)                       -0.060495   0.006274  -9.643  < 2e-16 ***
## sex                               0.069384   0.012121   5.724 1.05e-08 ***
## Scanner_Site11026                -0.005086   0.015356  -0.331   0.7405    
## Scanner_Site11027                 0.025424   0.011880   2.140   0.0323 *  
## ---
## Signif. codes:  0 '***' 0.001 '**' 0.01 '*' 0.05 '.' 0.1 ' ' 1
## 
## Residual standard error: 0.9878 on 38557 degrees of freedom
##   (1464 observations deleted due to missingness)
## Multiple R-squared:  0.02449,    Adjusted R-squared:  0.02434 
## F-statistic: 161.3 on 6 and 38557 DF,  p-value: < 2.2e-16
```

```
Beta1 <- 0.069384 
SE1 <- 0.012121
Beta2 <- 0.069384 
SE2 <- 0.012121

Z = ( Beta1 - Beta2) / sqrt((SE1^2)+ (SE2^2))
Z
```

```
## [1] 0
```

### III - Global Measures

```
## Load Data
DT<- fread("/scratch2/ukbio/biobank/Asymmetry/Asymmetry_2/data/Global_Measures_AI.csv")

## Add Scanner 
scanner_data <-fread("/scratch2/ukbio/biobank/Allometry/data/site_scanner_df.csv") 

scanner_data <- as.data.frame(scanner_data)
Merged_DF <- left_join(DT, scanner_data)
```

```
## Joining, by = "eid"
```

```
Merged_DF$Scanner_Site <- as.factor(Merged_DF$uk_biobank_assessment_centre_f54_2_0)
table(Merged_DF$Scanner_Site)
```

```
## 
## 11025 11026 11027 
## 24971  5070  9987
```

```
DT <- Merged_DF
DT1 <- setDT(DT)

# 2.Other ####
SUM_names <- c("Cerebellum_GM_sum", "Cortical_TSA_sum", "Cortical_GMV_sum",  
               "Cortical_MCT_sum","Subseg_Sum_no_DC","Subseg_Sum", 
               "Cerebellum_WM_sum", "Cerebellum_GM_ASEG_sum","Cerebral_WM_sum",
               "Cerebral_GM_ASEG_sum")

# Cortical_GMV L-R TBV 
Non_Linear_Model <- gam(scale(Cortical_GMV_AI) ~ s(scale(Cortical_GMV_sum))+ s(scale(TBV_AI_regional))+ 
                   scale(TBV) + 
                   sex +
                   Scanner_Site, data = DT1)
summary(Non_Linear_Model)
```

```
## 
## Family: gaussian 
## Link function: identity 
## 
## Formula:
## scale(Cortical_GMV_AI) ~ s(scale(Cortical_GMV_sum)) + s(scale(TBV_AI_regional)) + 
##     scale(TBV) + sex + Scanner_Site
## 
## Parametric coefficients:
##                    Estimate Std. Error t value Pr(>|t|)    
## (Intercept)        0.024379   0.002724   8.950  < 2e-16 ***
## scale(TBV)         0.050589   0.005862   8.630  < 2e-16 ***
## sex               -0.086816   0.005147 -16.869  < 2e-16 ***
## Scanner_Site11026 -0.039975   0.006493  -6.157 7.49e-10 ***
## Scanner_Site11027 -0.066889   0.005024 -13.314  < 2e-16 ***
## ---
## Signif. codes:  0 '***' 0.001 '**' 0.01 '*' 0.05 '.' 0.1 ' ' 1
## 
## Approximate significance of smooth terms:
##                              edf Ref.df         F  p-value    
## s(scale(Cortical_GMV_sum)) 1.922  2.460     7.865 0.000152 ***
## s(scale(TBV_AI_regional))  8.782  8.986 20285.250  < 2e-16 ***
## ---
## Signif. codes:  0 '***' 0.001 '**' 0.01 '*' 0.05 '.' 0.1 ' ' 1
## 
## R-sq.(adj) =  0.825   Deviance explained = 82.5%
## GCV = 0.17507  Scale est. = 0.175     n = 38710
```

```
Linear_Model <- lm(scale(Cortical_GMV_AI) ~ scale(Cortical_GMV_sum)+ (scale(TBV_AI_regional))+ 
                          scale(TBV) + 
                          sex +
                          Scanner_Site, data = DT1)
summary(Linear_Model)
```

```
## 
## Call:
## lm(formula = scale(Cortical_GMV_AI) ~ scale(Cortical_GMV_sum) + 
##     (scale(TBV_AI_regional)) + scale(TBV) + sex + Scanner_Site, 
##     data = DT1)
## 
## Residuals:
##     Min      1Q  Median      3Q     Max 
## -3.9825 -0.2628 -0.0001  0.2621  6.8091 
## 
## Coefficients:
##                          Estimate Std. Error t value Pr(>|t|)    
## (Intercept)              0.024894   0.002766   8.999  < 2e-16 ***
## scale(Cortical_GMV_sum) -0.024208   0.005760  -4.203 2.64e-05 ***
## scale(TBV_AI_regional)   0.909085   0.002170 418.915  < 2e-16 ***
## scale(TBV)               0.049688   0.005939   8.367  < 2e-16 ***
## sex                     -0.089518   0.005198 -17.221  < 2e-16 ***
## Scanner_Site11026       -0.040421   0.006594  -6.130 8.87e-10 ***
## Scanner_Site11027       -0.068333   0.005102 -13.394  < 2e-16 ***
## ---
## Signif. codes:  0 '***' 0.001 '**' 0.01 '*' 0.05 '.' 0.1 ' ' 1
## 
## Residual standard error: 0.4249 on 38703 degrees of freedom
##   (1318 observations deleted due to missingness)
## Multiple R-squared:  0.8195, Adjusted R-squared:  0.8194 
## F-statistic: 2.928e+04 on 6 and 38703 DF,  p-value: < 2.2e-16
```

```
Beta1 <- -0.086816 
SE1 <- 0.005147
Beta2 <- -0.089518 
SE2 <- 0.005198

Z = ( Beta1 - Beta2) / sqrt((SE1^2)+ (SE2^2))
Z
```

```
## [1] 0.3693725
```

```
# Cerebral WMV L-R TBV
Non_Linear_Model <- gam(scale(Cerebral_WM_AI) ~ scale(Cerebral_WM_sum)+ s(scale(TBV_AI_regional)) + 
                   scale(TBV) + 
                   sex +
                   Scanner_Site, data = DT1)
summary(Non_Linear_Model)
```

```
## 
## Family: gaussian 
## Link function: identity 
## 
## Formula:
## scale(Cerebral_WM_AI) ~ scale(Cerebral_WM_sum) + s(scale(TBV_AI_regional)) + 
##     scale(TBV) + sex + Scanner_Site
## 
## Parametric coefficients:
##                         Estimate Std. Error t value Pr(>|t|)    
## (Intercept)            -0.045526   0.005746  -7.924 2.37e-15 ***
## scale(Cerebral_WM_sum) -0.049283   0.014217  -3.466 0.000528 ***
## scale(TBV)              0.076865   0.014714   5.224 1.76e-07 ***
## sex                     0.122763   0.010857  11.307  < 2e-16 ***
## Scanner_Site11026       0.091495   0.013687   6.685 2.35e-11 ***
## Scanner_Site11027       0.110050   0.010590  10.392  < 2e-16 ***
## ---
## Signif. codes:  0 '***' 0.001 '**' 0.01 '*' 0.05 '.' 0.1 ' ' 1
## 
## Approximate significance of smooth terms:
##                             edf Ref.df    F p-value    
## s(scale(TBV_AI_regional)) 8.883  8.996 1130  <2e-16 ***
## ---
## Signif. codes:  0 '***' 0.001 '**' 0.01 '*' 0.05 '.' 0.1 ' ' 1
## 
## R-sq.(adj) =  0.217   Deviance explained = 21.8%
## GCV = 0.77789  Scale est. = 0.77759   n = 38710
```

```
Linear_Model <- lm(scale(Cerebral_WM_AI) ~ scale(Cerebral_WM_sum)+ (scale(TBV_AI_regional)) + 
                          scale(TBV) + 
                          sex +
                          Scanner_Site, data = DT1)
summary(Linear_Model)
```

```
## 
## Call:
## lm(formula = scale(Cerebral_WM_AI) ~ scale(Cerebral_WM_sum) + 
##     (scale(TBV_AI_regional)) + scale(TBV) + sex + Scanner_Site, 
##     data = DT1)
## 
## Residuals:
##      Min       1Q   Median       3Q      Max 
## -15.4801  -0.5417   0.0017   0.5475  10.0312 
## 
## Coefficients:
##                         Estimate Std. Error t value Pr(>|t|)    
## (Intercept)            -0.046482   0.005852  -7.943 2.03e-15 ***
## scale(Cerebral_WM_sum) -0.049788   0.014455  -3.444 0.000573 ***
## scale(TBV_AI_regional)  0.420189   0.004588  91.590  < 2e-16 ***
## scale(TBV)              0.080412   0.014955   5.377 7.63e-08 ***
## sex                     0.124416   0.011009  11.301  < 2e-16 ***
## Scanner_Site11026       0.093272   0.013941   6.691 2.25e-11 ***
## Scanner_Site11027       0.112560   0.010785  10.437  < 2e-16 ***
## ---
## Signif. codes:  0 '***' 0.001 '**' 0.01 '*' 0.05 '.' 0.1 ' ' 1
## 
## Residual standard error: 0.8982 on 38703 degrees of freedom
##   (1318 observations deleted due to missingness)
## Multiple R-squared:  0.1881, Adjusted R-squared:  0.188 
## F-statistic:  1494 on 6 and 38703 DF,  p-value: < 2.2e-16
```

```
Beta1 <- 0.122763 
SE1 <- 0.010857
Beta2 <- 0.124416 
SE2 <- 0.011009

Z = ( Beta1 - Beta2) / sqrt((SE1^2)+ (SE2^2))
Z
```

```
## [1] -0.1069075
```
