## Supplementary material for "Comparing Brain Asymmetries Independently of Brain Size": File_S5_Q4_Age_and_Sex_Interaction_Plots.html

Sex\_by\_Age\_Interactions


### Sex\_by\_Age\_Interactions

#### Significant Sex:Age Interactions

p < 0.05 / (5\*306) second graph for each measure is a zoom of the first graph to better visualize the interaction. GMV: Grey Matter Volume

```
## `geom_smooth()` using method = 'gam' and formula 'y ~ s(x, bs = "cs")'
```

```
## `geom_smooth()` using method = 'gam' and formula 'y ~ s(x, bs = "cs")'
```

```
## `geom_smooth()` using method = 'gam' and formula 'y ~ s(x, bs = "cs")'
```

```
## `geom_smooth()` using method = 'gam' and formula 'y ~ s(x, bs = "cs")'
```

```
## `geom_smooth()` using method = 'gam' and formula 'y ~ s(x, bs = "cs")'
```

```
## `geom_smooth()` using method = 'gam' and formula 'y ~ s(x, bs = "cs")'
```

```
## `geom_smooth()` using method = 'gam' and formula 'y ~ s(x, bs = "cs")'
```

```
## `geom_smooth()` using method = 'gam' and formula 'y ~ s(x, bs = "cs")'
```

```
## `geom_smooth()` using method = 'gam' and formula 'y ~ s(x, bs = "cs")'
```

```
## `geom_smooth()` using method = 'gam' and formula 'y ~ s(x, bs = "cs")'
```

```
## `geom_smooth()` using method = 'gam' and formula 'y ~ s(x, bs = "cs")'
```

```
## `geom_smooth()` using method = 'gam' and formula 'y ~ s(x, bs = "cs")'
```

```
## `geom_smooth()` using method = 'gam' and formula 'y ~ s(x, bs = "cs")'
```

```
## `geom_smooth()` using method = 'gam' and formula 'y ~ s(x, bs = "cs")'
```

```
## `geom_smooth()` using method = 'gam' and formula 'y ~ s(x, bs = "cs")'
```

```
## `geom_smooth()` using method = 'gam' and formula 'y ~ s(x, bs = "cs")'
```

```
## `geom_smooth()` using method = 'gam' and formula 'y ~ s(x, bs = "cs")'
```

```
## `geom_smooth()` using method = 'gam' and formula 'y ~ s(x, bs = "cs")'
```
