## Supplementary material for "Comparing Brain Asymmetries Independently of Brain Size": File_S6_Exploratory_Analyses_Asymmetry_Significance_CohensD.html

Histograms 1 Sample T Test


### Histograms 1 Sample T Test

### 1. Destrieux

L: Left Hemisphere R: Right Hemisphere

```
## 
## Not Significant     Significant 
##               1              73
```

```
## 
## Not Significant     Significant 
##               4              70
```

```
## 
## Not Significant     Significant 
##               1              73
```

### 2. Desikan

L: Left Hemisphere R: Right Hemisphere

```
## 
## Significant 
##          31
```

```
## 
## Not Significant     Significant 
##               2              29
```

```
## 
## Significant 
##          31
```
