## Supplementary material for "Comparing Brain Asymmetries Independently of Brain Size": File_S7_GGseg_Sex_Age_Graphs_SA_MT_DK_Williams_Kong_All.html

Desikan Sex Effects


### Desikan Sex Effects

### I - DKT and DK

#### A) Sex

##### 1. Mean Thickness

###### a. Williams no L+R and Global AI

```
## merging atlas and data by 'atlas', 'type', 'hemi', 'side', 'roi'
```

###### b. Kong no L+R and Global AI

```
## merging atlas and data by 'atlas', 'type', 'hemi', 'side', 'roi'
```

```
## Warning: Some data not merged properly. Check for naming errors in data:
##   atlas type  hemi  side  region label roi   Sex_Estimate Region     pval Study 
##   <chr> <chr> <chr> <chr> <chr>  <chr> <chr>        <dbl> <chr>     <dbl> <chr> 
## 1 <NA>  <NA>  <NA>  <NA>  <NA>   <NA>  <NA>      -0.00438 bankssts  0.814 DK_Ko…
## 2 <NA>  <NA>  <NA>  <NA>  <NA>   <NA>  <NA>       0.0241  frontalp… 0.276 DK_Ko…
## 3 <NA>  <NA>  <NA>  <NA>  <NA>   <NA>  <NA>      -0.0246  temporal… 0.234 DK_Ko…
## # … with 4 more variables: region.x <chr>, AI <dbl>, region.y <chr>,
## #   geometry <MULTIPOLYGON>
```

###### c. Williams with L+R and Global AI

```
## merging atlas and data by 'atlas', 'type', 'hemi', 'side', 'roi'
```

###### d. Williams with TBV

```
## merging atlas and data by 'atlas', 'type', 'hemi', 'side', 'roi'
```

##### 2. Surface Areas

###### a. Williams no L+R and Global AI

```
## merging atlas and data by 'atlas', 'type', 'hemi', 'side', 'roi'
```

###### b. Kong no L+R and Global AI

```
## merging atlas and data by 'atlas', 'type', 'hemi', 'side', 'roi'
```

```
## Warning: Some data not merged properly. Check for naming errors in data:
##   atlas type  hemi  side  region label roi   Sex_Estimate Region      pval Study
##   <chr> <chr> <chr> <chr> <chr>  <chr> <chr>        <dbl> <chr>      <dbl> <chr>
## 1 <NA>  <NA>  <NA>  <NA>  <NA>   <NA>  <NA>      -0.00980 bankssts 5.65e-1 DK_K…
## 2 <NA>  <NA>  <NA>  <NA>  <NA>   <NA>  <NA>      -0.0406  frontal… 1.31e-2 DK_K…
## 3 <NA>  <NA>  <NA>  <NA>  <NA>   <NA>  <NA>       0.0638  tempora… 5.15e-4 DK_K…
## # … with 4 more variables: region.x <chr>, AI <dbl>, region.y <chr>,
## #   geometry <MULTIPOLYGON>
```

###### c. Williams with L+R and Global AI

```
## merging atlas and data by 'atlas', 'type', 'hemi', 'side', 'roi'
```

###### d. Williams with TBV

```
## merging atlas and data by 'atlas', 'type', 'hemi', 'side', 'roi'
```

#### B) Age

##### 1. Mean Thickness

###### a. Williams no L+R and Global AI

```
## merging atlas and data by 'atlas', 'type', 'hemi', 'side', 'roi'
```

###### b. Kong no L+R and Global AI

```
## merging atlas and data by 'atlas', 'type', 'hemi', 'side', 'roi'
```

```
## Warning: Some data not merged properly. Check for naming errors in data:
##   atlas type  hemi  side  region label roi   Age_Estimate Region     pval Study 
##   <chr> <chr> <chr> <chr> <chr>  <chr> <chr>        <dbl> <chr>     <dbl> <chr> 
## 1 <NA>  <NA>  <NA>  <NA>  <NA>   <NA>  <NA>       0.00140 bankssts  0.866 DK_Ko…
## 2 <NA>  <NA>  <NA>  <NA>  <NA>   <NA>  <NA>       0.00774 frontalp… 0.410 DK_Ko…
## 3 <NA>  <NA>  <NA>  <NA>  <NA>   <NA>  <NA>      -0.00993 temporal… 0.221 DK_Ko…
## # … with 4 more variables: region.x <chr>, AI <dbl>, region.y <chr>,
## #   geometry <MULTIPOLYGON>
```

###### c. Williams with L+R and Global AI

```
## merging atlas and data by 'atlas', 'type', 'hemi', 'side', 'roi'
```

###### d. Williams with TBV

```
## merging atlas and data by 'atlas', 'type', 'hemi', 'side', 'roi'
```

##### 2. Surface Areas

###### a. Williams no L+R and Global AI

```
## merging atlas and data by 'atlas', 'type', 'hemi', 'side', 'roi'
```

###### b. Kong no L+R and Global AI

```
## merging atlas and data by 'atlas', 'type', 'hemi', 'side', 'roi'
```

```
## Warning: Some data not merged properly. Check for naming errors in data:
##   atlas type  hemi  side  region label roi   Age_Estimate Region      pval Study
##   <chr> <chr> <chr> <chr> <chr>  <chr> <chr>        <dbl> <chr>      <dbl> <chr>
## 1 <NA>  <NA>  <NA>  <NA>  <NA>   <NA>  <NA>      -0.0164  bankssts  0.0492 DK_K…
## 2 <NA>  <NA>  <NA>  <NA>  <NA>   <NA>  <NA>      -0.00763 frontalp… 0.401  DK_K…
## 3 <NA>  <NA>  <NA>  <NA>  <NA>   <NA>  <NA>       0.0120  temporal… 0.196  DK_K…
## # … with 4 more variables: region.x <chr>, AI <dbl>, region.y <chr>,
## #   geometry <MULTIPOLYGON>
```

###### c. Williams with L+R and Global AI

```
## merging atlas and data by 'atlas', 'type', 'hemi', 'side', 'roi'
```

###### d. Williams with TBV

```
## merging atlas and data by 'atlas', 'type', 'hemi', 'side', 'roi'
```

### II. Destrieux

```
## # A tibble: 11 × 6
##    repo                       ggseg ggseg3d source comment             package  
##    <chr>                      <lgl> <lgl>   <chr>  <chr>               <chr>    
##  1 LCBC-UiO/ggsegYeo2011      TRUE  TRUE    github both 17 and 7 Netw… ggsegYeo…
##  2 LCBC-UiO/ggsegDesterieux   FALSE TRUE    github the 2009 atlas      ggsegDes…
##  3 LCBC-UiO/ggsegChen         FALSE TRUE    github both thickness and… ggsegChen
##  4 LCBC-UiO/ggsegSchaefer     FALSE TRUE    github both 17 and 7 netw… ggsegSch…
##  5 LCBC-UiO/ggsegGlasser      TRUE  TRUE    github full atlas          ggsegGla…
##  6 LCBC-UiO/ggsegJHU          TRUE  TRUE    github white tract atlas   ggsegJHU 
##  7 LCBC-UiO/ggsegTracula      TRUE  TRUE    github white tract atlas   ggsegTra…
##  8 LCBC-UiO/ggsegICBM         FALSE TRUE    github white tract atlas   ggsegICBM
##  9 LCBC-UiO/ggsegHO           TRUE  FALSE   github Harvard-Oxford cor… ggsegHO  
## 10 LCBC-UiO/ggsegDefaultExtra TRUE  FALSE   github extra 2d view for … ggsegDef…
## 11 LCBC-UiO/ggsegDKT          TRUE  TRUE    github Desikan-Killiany-T… ggsegDKT
```

```
##   [1] "volume_of_lateralventricle_AI_old"                        
##   [2] "volume_of_inflatvent_AI_old"                              
##   [3] "volume_of_caudate_AI_old"                                 
##   [4] "volume_of_putamen_AI_old"                                 
##   [5] "volume_of_pallidum_AI_old"                                
##   [6] "volume_of_accumbensarea_AI_old"                           
##   [7] "volume_of_ventraldc_AI_old"                               
##   [8] "volume_of_choroidplexus_AI_old"                           
##   [9] "volume_of_grey_matter_in_iiv_cerebellum_AI_old_AI_old"    
##  [10] "volume_of_grey_matter_in_v_cerebellum_AI_old_AI_old"      
##  [11] "volume_of_grey_matter_in_vi_cerebellum_AI_old_AI_old"     
##  [12] "volume_of_grey_matter_in_crus_i_cerebellum_AI_old_AI_old" 
##  [13] "volume_of_grey_matter_in_crus_ii_cerebellum_AI_old_AI_old"
##  [14] "volume_of_grey_matter_in_viib_cerebellum_AI_old_AI_old"   
##  [15] "volume_of_grey_matter_in_viiia_cerebellum_AI_old_AI_old"  
##  [16] "volume_of_grey_matter_in_viiib_cerebellum_AI_old_AI_old"  
##  [17] "volume_of_grey_matter_in_ix_cerebellum_AI_old_AI_old"     
##  [18] "volume_of_grey_matter_in_x_cerebellum_AI_old_AI_old"      
##  [19] "volume_of_lateralnucleus_AI_old"                          
##  [20] "volume_of_basalnucleus_AI_old"                            
##  [21] "volume_of_accessorybasalnucleus_AI_old"                   
##  [22] "volume_of_anterioramygdaloidareaaaa_AI_old"               
##  [23] "volume_of_centralnucleus_AI_old"                          
##  [24] "volume_of_medialnucleus_AI_old"                           
##  [25] "volume_of_corticalnucleus_AI_old"                         
##  [26] "volume_of_corticoamygdaloidtransitio_AI_old"              
##  [27] "volume_of_paralaminarnucleus_AI_old"                      
##  [28] "volume_of_wholeamygdala_AI_old"                           
##  [29] "volume_of_hippocampaltail_AI_old"                         
##  [30] "volume_of_subiculumbody_AI_old"                           
##  [31] "volume_of_ca1body_AI_old"                                 
##  [32] "volume_of_subiculumhead_AI_old"                           
##  [33] "volume_of_hippocampalfissure_AI_old"                      
##  [34] "volume_of_presubiculumhead_AI_old"                        
##  [35] "volume_of_ca1head_AI_old"                                 
##  [36] "volume_of_presubiculumbody_AI_old"                        
##  [37] "volume_of_parasubiculum_AI_old"                           
##  [38] "volume_of_molecularlayerhphead_AI_old"                    
##  [39] "volume_of_molecularlayerhpbody_AI_old"                    
##  [40] "volume_of_gcmldghead_AI_old"                              
##  [41] "volume_of_ca3body_AI_old"                                 
##  [42] "volume_of_gcmldgbody_AI_old"                              
##  [43] "volume_of_ca4head_AI_old"                                 
##  [44] "volume_of_ca4body_AI_old"                                 
##  [45] "volume_of_fimbria_AI_old"                                 
##  [46] "volume_of_ca3head_AI_old"                                 
##  [47] "volume_of_hata_AI_old"                                    
##  [48] "volume_of_wholehippocampus_AI_old"                        
##  [49] "volume_of_mgn_AI_old"                                     
##  [50] "volume_of_lgn_AI_old"                                     
##  [51] "volume_of_pui_AI_old"                                     
##  [52] "volume_of_pum_AI_old"                                     
##  [53] "volume_of_lsg_AI_old"                                     
##  [54] "volume_of_vpl_AI_old"                                     
##  [55] "volume_of_cm_AI_old"                                      
##  [56] "volume_of_vla_AI_old"                                     
##  [57] "volume_of_pua_AI_old"                                     
##  [58] "volume_of_mdm_AI_old"                                     
##  [59] "volume_of_pf_AI_old"                                      
##  [60] "volume_of_vamc_AI_old"                                    
##  [61] "volume_of_mdl_AI_old"                                     
##  [62] "volume_of_cem_AI_old"                                     
##  [63] "volume_of_va_AI_old"                                      
##  [64] "volume_of_mvre_AI_old"                                    
##  [65] "volume_of_vm_AI_old"                                      
##  [66] "volume_of_cl_AI_old"                                      
##  [67] "volume_of_pul_AI_old"                                     
##  [68] "volume_of_pt_AI_old"                                      
##  [69] "volume_of_av_AI_old"                                      
##  [70] "volume_of_pc_AI_old"                                      
##  [71] "volume_of_vlp_AI_old"                                     
##  [72] "volume_of_lp_AI_old"                                      
##  [73] "volume_of_ld_AI_old"                                      
##  [74] "volume_of_wholethalamus_AI_old"                           
##  [75] "volume_of_gsfrontomargin_AI_old"                          
##  [76] "volume_of_gsoccipitalinf_AI_old"                          
##  [77] "volume_of_gsparacentral_AI_old"                           
##  [78] "volume_of_gssubcentral_AI_old"                            
##  [79] "volume_of_gstransvfrontopol_AI_old"                       
##  [80] "volume_of_gscingulant_AI_old"                             
##  [81] "volume_of_gscingulmidant_AI_old"                          
##  [82] "volume_of_gscingulmidpost_AI_old"                         
##  [83] "volume_of_gcingulpostdorsal_AI_old"                       
##  [84] "volume_of_gcingulpostventral_AI_old"                      
##  [85] "volume_of_gcuneus_AI_old"                                 
##  [86] "volume_of_gfrontinfopercular_AI_old"                      
##  [87] "volume_of_gfrontinforbital_AI_old"                        
##  [88] "volume_of_gfrontinftriangul_AI_old"                       
##  [89] "volume_of_gfrontmiddle_AI_old"                            
##  [90] "volume_of_gfrontsup_AI_old"                               
##  [91] "volume_of_ginslgscentins_AI_old"                          
##  [92] "volume_of_ginsularshort_AI_old"                           
##  [93] "volume_of_goccipitalmiddle_AI_old"                        
##  [94] "volume_of_goccipitalsup_AI_old"                           
##  [95] "volume_of_goctemplatfusifor_AI_old"                       
##  [96] "volume_of_goctempmedlingual_AI_old"                       
##  [97] "volume_of_goctempmedparahip_AI_old"                       
##  [98] "volume_of_gorbital_AI_old"                                
##  [99] "volume_of_gparietinfangular_AI_old"                       
## [100] "volume_of_gparietinfsupramar_AI_old"                      
## [101] "volume_of_gparietalsup_AI_old"                            
## [102] "volume_of_gpostcentral_AI_old"                            
## [103] "volume_of_gprecentral_AI_old"                             
## [104] "volume_of_gprecuneus_AI_old"                              
## [105] "volume_of_grectus_AI_old"                                 
## [106] "volume_of_gsubcallosal_AI_old"                            
## [107] "volume_of_gtempsupgttransv_AI_old"                        
## [108] "volume_of_gtempsuplateral_AI_old"                         
## [109] "volume_of_gtempsupplanpolar_AI_old"                       
## [110] "volume_of_gtempsupplantempo_AI_old"                       
## [111] "volume_of_gtemporalinf_AI_old"                            
## [112] "volume_of_gtemporalmiddle_AI_old"                         
## [113] "volume_of_latfisanthorizont_AI_old"                       
## [114] "volume_of_latfisantvertical_AI_old"                       
## [115] "volume_of_latfispost_AI_old"                              
## [116] "volume_of_poleoccipital_AI_old"                           
## [117] "volume_of_poletemporal_AI_old"                            
## [118] "volume_of_scalcarine_AI_old"                              
## [119] "volume_of_scentral_AI_old"                                
## [120] "volume_of_scingulmarginalis_AI_old"                       
## [121] "volume_of_scircularinsulaant_AI_old"                      
## [122] "volume_of_scircularinsulainf_AI_old"                      
## [123] "volume_of_scircularinsulasup_AI_old"                      
## [124] "volume_of_scollattransvant_AI_old"                        
## [125] "volume_of_scollattransvpost_AI_old"                       
## [126] "volume_of_sfrontinf_AI_old"                               
## [127] "volume_of_sfrontmiddle_AI_old"                            
## [128] "volume_of_sfrontsup_AI_old"                               
## [129] "volume_of_sintermprimjensen_AI_old"                       
## [130] "volume_of_sintraparietptrans_AI_old"                      
## [131] "volume_of_socmiddlelunatus_AI_old"                        
## [132] "volume_of_socsuptransversal_AI_old"                       
## [133] "volume_of_soccipitalant_AI_old"                           
## [134] "volume_of_soctemplat_AI_old"                              
## [135] "volume_of_soctempmedlingual_AI_old"                       
## [136] "volume_of_sorbitallateral_AI_old"                         
## [137] "volume_of_sorbitalmedolfact_AI_old"                       
## [138] "volume_of_sorbitalhshaped_AI_old"                         
## [139] "volume_of_sparietooccipital_AI_old"                       
## [140] "volume_of_spericallosal_AI_old"                           
## [141] "volume_of_spostcentral_AI_old"                            
## [142] "volume_of_sprecentralinfpart_AI_old"                      
## [143] "volume_of_sprecentralsuppart_AI_old"                      
## [144] "volume_of_ssuborbital_AI_old"                             
## [145] "volume_of_ssubparietal_AI_old"                            
## [146] "volume_of_stemporalinf_AI_old"                            
## [147] "volume_of_stemporalsup_AI_old"                            
## [148] "volume_of_stemporaltransverse_AI_old"
```

```
##  [1] "volume_of_gsfrontomargin_AI_old"     
##  [2] "volume_of_gsoccipitalinf_AI_old"     
##  [3] "volume_of_gsparacentral_AI_old"      
##  [4] "volume_of_gssubcentral_AI_old"       
##  [5] "volume_of_gstransvfrontopol_AI_old"  
##  [6] "volume_of_gscingulant_AI_old"        
##  [7] "volume_of_gscingulmidant_AI_old"     
##  [8] "volume_of_gscingulmidpost_AI_old"    
##  [9] "volume_of_gcingulpostdorsal_AI_old"  
## [10] "volume_of_gcingulpostventral_AI_old" 
## [11] "volume_of_gcuneus_AI_old"            
## [12] "volume_of_gfrontinfopercular_AI_old" 
## [13] "volume_of_gfrontinforbital_AI_old"   
## [14] "volume_of_gfrontinftriangul_AI_old"  
## [15] "volume_of_gfrontmiddle_AI_old"       
## [16] "volume_of_gfrontsup_AI_old"          
## [17] "volume_of_ginslgscentins_AI_old"     
## [18] "volume_of_ginsularshort_AI_old"      
## [19] "volume_of_goccipitalmiddle_AI_old"   
## [20] "volume_of_goccipitalsup_AI_old"      
## [21] "volume_of_goctemplatfusifor_AI_old"  
## [22] "volume_of_goctempmedlingual_AI_old"  
## [23] "volume_of_goctempmedparahip_AI_old"  
## [24] "volume_of_gorbital_AI_old"           
## [25] "volume_of_gparietinfangular_AI_old"  
## [26] "volume_of_gparietinfsupramar_AI_old" 
## [27] "volume_of_gparietalsup_AI_old"       
## [28] "volume_of_gpostcentral_AI_old"       
## [29] "volume_of_gprecentral_AI_old"        
## [30] "volume_of_gprecuneus_AI_old"         
## [31] "volume_of_grectus_AI_old"            
## [32] "volume_of_gsubcallosal_AI_old"       
## [33] "volume_of_gtempsupgttransv_AI_old"   
## [34] "volume_of_gtempsuplateral_AI_old"    
## [35] "volume_of_gtempsupplanpolar_AI_old"  
## [36] "volume_of_gtempsupplantempo_AI_old"  
## [37] "volume_of_gtemporalinf_AI_old"       
## [38] "volume_of_gtemporalmiddle_AI_old"    
## [39] "volume_of_latfisanthorizont_AI_old"  
## [40] "volume_of_latfisantvertical_AI_old"  
## [41] "volume_of_latfispost_AI_old"         
## [42] "volume_of_poleoccipital_AI_old"      
## [43] "volume_of_poletemporal_AI_old"       
## [44] "volume_of_scalcarine_AI_old"         
## [45] "volume_of_scentral_AI_old"           
## [46] "volume_of_scingulmarginalis_AI_old"  
## [47] "volume_of_scircularinsulaant_AI_old" 
## [48] "volume_of_scircularinsulainf_AI_old" 
## [49] "volume_of_scircularinsulasup_AI_old" 
## [50] "volume_of_scollattransvant_AI_old"   
## [51] "volume_of_scollattransvpost_AI_old"  
## [52] "volume_of_sfrontinf_AI_old"          
## [53] "volume_of_sfrontmiddle_AI_old"       
## [54] "volume_of_sfrontsup_AI_old"          
## [55] "volume_of_sintermprimjensen_AI_old"  
## [56] "volume_of_sintraparietptrans_AI_old" 
## [57] "volume_of_socmiddlelunatus_AI_old"   
## [58] "volume_of_socsuptransversal_AI_old"  
## [59] "volume_of_soccipitalant_AI_old"      
## [60] "volume_of_soctemplat_AI_old"         
## [61] "volume_of_soctempmedlingual_AI_old"  
## [62] "volume_of_sorbitallateral_AI_old"    
## [63] "volume_of_sorbitalmedolfact_AI_old"  
## [64] "volume_of_sorbitalhshaped_AI_old"    
## [65] "volume_of_sparietooccipital_AI_old"  
## [66] "volume_of_spericallosal_AI_old"      
## [67] "volume_of_spostcentral_AI_old"       
## [68] "volume_of_sprecentralinfpart_AI_old" 
## [69] "volume_of_sprecentralsuppart_AI_old" 
## [70] "volume_of_ssuborbital_AI_old"        
## [71] "volume_of_ssubparietal_AI_old"       
## [72] "volume_of_stemporalinf_AI_old"       
## [73] "volume_of_stemporalsup_AI_old"       
## [74] "volume_of_stemporaltransverse_AI_old"
```

```
## merging atlas and data by 'atlas', 'type', 'hemi', 'side', 'region', 'roi'
```

```
## merging atlas and data by 'atlas', 'type', 'hemi', 'side', 'region', 'roi'
```

```
##  [1] "area_of_gsfrontomargin_AI_old"      "area_of_gsoccipitalinf_AI_old"     
##  [3] "area_of_gsparacentral_AI_old"       "area_of_gssubcentral_AI_old"       
##  [5] "area_of_gstransvfrontopol_AI_old"   "area_of_gscingulant_AI_old"        
##  [7] "area_of_gscingulmidant_AI_old"      "area_of_gscingulmidpost_AI_old"    
##  [9] "area_of_gcingulpostdorsal_AI_old"   "area_of_gcingulpostventral_AI_old" 
## [11] "area_of_gcuneus_AI_old"             "area_of_gfrontinfopercular_AI_old" 
## [13] "area_of_gfrontinforbital_AI_old"    "area_of_gfrontinftriangul_AI_old"  
## [15] "area_of_gfrontmiddle_AI_old"        "area_of_gfrontsup_AI_old"          
## [17] "area_of_ginslgscentins_AI_old"      "area_of_ginsularshort_AI_old"      
## [19] "area_of_goccipitalmiddle_AI_old"    "area_of_goccipitalsup_AI_old"      
## [21] "area_of_goctemplatfusifor_AI_old"   "area_of_goctempmedlingual_AI_old"  
## [23] "area_of_goctempmedparahip_AI_old"   "area_of_gorbital_AI_old"           
## [25] "area_of_gparietinfangular_AI_old"   "area_of_gparietinfsupramar_AI_old" 
## [27] "area_of_gparietalsup_AI_old"        "area_of_gpostcentral_AI_old"       
## [29] "area_of_gprecentral_AI_old"         "area_of_gprecuneus_AI_old"         
## [31] "area_of_grectus_AI_old"             "area_of_gsubcallosal_AI_old"       
## [33] "area_of_gtempsupgttransv_AI_old"    "area_of_gtempsuplateral_AI_old"    
## [35] "area_of_gtempsupplanpolar_AI_old"   "area_of_gtempsupplantempo_AI_old"  
## [37] "area_of_gtemporalinf_AI_old"        "area_of_gtemporalmiddle_AI_old"    
## [39] "area_of_latfisanthorizont_AI_old"   "area_of_latfisantvertical_AI_old"  
## [41] "area_of_latfispost_AI_old"          "area_of_poleoccipital_AI_old"      
## [43] "area_of_poletemporal_AI_old"        "area_of_scalcarine_AI_old"         
## [45] "area_of_scentral_AI_old"            "area_of_scingulmarginalis_AI_old"  
## [47] "area_of_scircularinsulaant_AI_old"  "area_of_scircularinsulainf_AI_old" 
## [49] "area_of_scircularinsulasup_AI_old"  "area_of_scollattransvant_AI_old"   
## [51] "area_of_scollattransvpost_AI_old"   "area_of_sfrontinf_AI_old"          
## [53] "area_of_sfrontmiddle_AI_old"        "area_of_sfrontsup_AI_old"          
## [55] "area_of_sintermprimjensen_AI_old"   "area_of_sintraparietptrans_AI_old" 
## [57] "area_of_socmiddlelunatus_AI_old"    "area_of_socsuptransversal_AI_old"  
## [59] "area_of_soccipitalant_AI_old"       "area_of_soctemplat_AI_old"         
## [61] "area_of_soctempmedlingual_AI_old"   "area_of_sorbitallateral_AI_old"    
## [63] "area_of_sorbitalmedolfact_AI_old"   "area_of_sorbitalhshaped_AI_old"    
## [65] "area_of_sparietooccipital_AI_old"   "area_of_spericallosal_AI_old"      
## [67] "area_of_spostcentral_AI_old"        "area_of_sprecentralinfpart_AI_old" 
## [69] "area_of_sprecentralsuppart_AI_old"  "area_of_ssuborbital_AI_old"        
## [71] "area_of_ssubparietal_AI_old"        "area_of_stemporalinf_AI_old"       
## [73] "area_of_stemporalsup_AI_old"        "area_of_stemporaltransverse_AI_old"
```

```
## merging atlas and data by 'atlas', 'type', 'hemi', 'side', 'region', 'roi'
```

```
## merging atlas and data by 'atlas', 'type', 'hemi', 'side', 'region', 'roi'
```

```
##  [1] "mean_thickness_of_gsfrontomargin_AI_old"     
##  [2] "mean_thickness_of_gsoccipitalinf_AI_old"     
##  [3] "mean_thickness_of_gsparacentral_AI_old"      
##  [4] "mean_thickness_of_gssubcentral_AI_old"       
##  [5] "mean_thickness_of_gstransvfrontopol_AI_old"  
##  [6] "mean_thickness_of_gscingulant_AI_old"        
##  [7] "mean_thickness_of_gscingulmidant_AI_old"     
##  [8] "mean_thickness_of_gscingulmidpost_AI_old"    
##  [9] "mean_thickness_of_gcingulpostdorsal_AI_old"  
## [10] "mean_thickness_of_gcingulpostventral_AI_old" 
## [11] "mean_thickness_of_gcuneus_AI_old"            
## [12] "mean_thickness_of_gfrontinfopercular_AI_old" 
## [13] "mean_thickness_of_gfrontinforbital_AI_old"   
## [14] "mean_thickness_of_gfrontinftriangul_AI_old"  
## [15] "mean_thickness_of_gfrontmiddle_AI_old"       
## [16] "mean_thickness_of_gfrontsup_AI_old"          
## [17] "mean_thickness_of_ginslgscentins_AI_old"     
## [18] "mean_thickness_of_ginsularshort_AI_old"      
## [19] "mean_thickness_of_goccipitalmiddle_AI_old"   
## [20] "mean_thickness_of_goccipitalsup_AI_old"      
## [21] "mean_thickness_of_goctemplatfusifor_AI_old"  
## [22] "mean_thickness_of_goctempmedlingual_AI_old"  
## [23] "mean_thickness_of_goctempmedparahip_AI_old"  
## [24] "mean_thickness_of_gorbital_AI_old"           
## [25] "mean_thickness_of_gparietinfangular_AI_old"  
## [26] "mean_thickness_of_gparietinfsupramar_AI_old" 
## [27] "mean_thickness_of_gparietalsup_AI_old"       
## [28] "mean_thickness_of_gpostcentral_AI_old"       
## [29] "mean_thickness_of_gprecentral_AI_old"        
## [30] "mean_thickness_of_gprecuneus_AI_old"         
## [31] "mean_thickness_of_grectus_AI_old"            
## [32] "mean_thickness_of_gsubcallosal_AI_old"       
## [33] "mean_thickness_of_gtempsupgttransv_AI_old"   
## [34] "mean_thickness_of_gtempsuplateral_AI_old"    
## [35] "mean_thickness_of_gtempsupplanpolar_AI_old"  
## [36] "mean_thickness_of_gtempsupplantempo_AI_old"  
## [37] "mean_thickness_of_gtemporalinf_AI_old"       
## [38] "mean_thickness_of_gtemporalmiddle_AI_old"    
## [39] "mean_thickness_of_latfisanthorizont_AI_old"  
## [40] "mean_thickness_of_latfisantvertical_AI_old"  
## [41] "mean_thickness_of_latfispost_AI_old"         
## [42] "mean_thickness_of_poleoccipital_AI_old"      
## [43] "mean_thickness_of_poletemporal_AI_old"       
## [44] "mean_thickness_of_scalcarine_AI_old"         
## [45] "mean_thickness_of_scentral_AI_old"           
## [46] "mean_thickness_of_scingulmarginalis_AI_old"  
## [47] "mean_thickness_of_scircularinsulaant_AI_old" 
## [48] "mean_thickness_of_scircularinsulainf_AI_old" 
## [49] "mean_thickness_of_scircularinsulasup_AI_old" 
## [50] "mean_thickness_of_scollattransvant_AI_old"   
## [51] "mean_thickness_of_scollattransvpost_AI_old"  
## [52] "mean_thickness_of_sfrontinf_AI_old"          
## [53] "mean_thickness_of_sfrontmiddle_AI_old"       
## [54] "mean_thickness_of_sfrontsup_AI_old"          
## [55] "mean_thickness_of_sintermprimjensen_AI_old"  
## [56] "mean_thickness_of_sintraparietptrans_AI_old" 
## [57] "mean_thickness_of_socmiddlelunatus_AI_old"   
## [58] "mean_thickness_of_socsuptransversal_AI_old"  
## [59] "mean_thickness_of_soccipitalant_AI_old"      
## [60] "mean_thickness_of_soctemplat_AI_old"         
## [61] "mean_thickness_of_soctempmedlingual_AI_old"  
## [62] "mean_thickness_of_sorbitallateral_AI_old"    
## [63] "mean_thickness_of_sorbitalmedolfact_AI_old"  
## [64] "mean_thickness_of_sorbitalhshaped_AI_old"    
## [65] "mean_thickness_of_sparietooccipital_AI_old"  
## [66] "mean_thickness_of_spericallosal_AI_old"      
## [67] "mean_thickness_of_spostcentral_AI_old"       
## [68] "mean_thickness_of_sprecentralinfpart_AI_old" 
## [69] "mean_thickness_of_sprecentralsuppart_AI_old" 
## [70] "mean_thickness_of_ssuborbital_AI_old"        
## [71] "mean_thickness_of_ssubparietal_AI_old"       
## [72] "mean_thickness_of_stemporalinf_AI_old"       
## [73] "mean_thickness_of_stemporalsup_AI_old"       
## [74] "mean_thickness_of_stemporaltransverse_AI_old"
```

```
## merging atlas and data by 'atlas', 'type', 'hemi', 'side', 'region', 'roi'
```

```
## merging atlas and data by 'atlas', 'type', 'hemi', 'side', 'region', 'roi'
```
