## Supplementary material for "Comparing Brain Asymmetries Independently of Brain Size": File_S8_Guadalupe_Williams_Subcortical_Correlation_AI.html

Guadalupe Subcortical Correlations: Guadalupe and Williams


### Guadalupe Subcortical Correlations: Guadalupe and Williams

## AI

```
## `geom_smooth()` using formula 'y ~ x'
```

#### Sex Effet

```
## `geom_smooth()` using formula 'y ~ x'
```

#### Age Effet

```
## `geom_smooth()` using formula 'y ~ x'
```
