## Supplementary material for "Comparing Brain Asymmetries Independently of Brain Size": File_S9_Sarica_Williams_Hippocampal_Subfield_Correlations.html

Sarica and Williams Hippocampal Subfeild Correlations


### Sarica and Williams Hippocampal Subfeild Correlations

```
## Warning: Removed 1 rows containing non-finite values (stat_cor).
```

```
## `geom_smooth()` using formula 'y ~ x'
```

```
## Warning: Removed 1 rows containing non-finite values (stat_smooth).
```

```
## Warning: Removed 1 rows containing missing values (geom_point).
```

```
## Warning: Removed 1 rows containing missing values (geom_label_repel).
```
