## Supplementary material for "Comparing Brain Asymmetries Independently of Brain Size": GGseg_Sex_Age_Graphs_SA_MT_DK_Williams_Kong_Sig.html

Desikan Sex Effects


### Desikan Sex Effects

### I - DKT and DK

#### A) Sex

##### 1. Mean Thickness

###### a. Williams no L+R and Global AI

```
## merging atlas and data by 'atlas', 'type', 'hemi', 'side', 'roi'
```

###### b. Kong no L+R and Global AI

```
## merging atlas and data by 'atlas', 'type', 'hemi', 'side', 'roi'
```

```
## Warning: Some data not merged properly. Check for naming errors in data:
##   atlas type  hemi  side  region label roi   Sex_Estimate1 Region     pval Study
##   <chr> <chr> <chr> <chr> <chr>  <chr> <chr>         <dbl> <chr>     <dbl> <chr>
## 1 <NA>  <NA>  <NA>  <NA>  <NA>   <NA>  <NA>       -0.00438 bankssts  0.814 DK_K…
## 2 <NA>  <NA>  <NA>  <NA>  <NA>   <NA>  <NA>        0.0241  frontalp… 0.276 DK_K…
## 3 <NA>  <NA>  <NA>  <NA>  <NA>   <NA>  <NA>       -0.0246  temporal… 0.234 DK_K…
## # … with 5 more variables: Sex_Estimate <dbl>, region.x <chr>, AI <dbl>,
## #   region.y <chr>, geometry <MULTIPOLYGON>
```

###### c. Williams with L+R and Global AI

```
##  [1]  0.0000  0.0000  0.0000  0.0000  0.0000  0.0814  0.0000 -0.0922  0.0000
## [10] -0.1438 -0.0539 -0.0696  0.0000 -0.0635  0.0596  0.0000 -0.0652  0.0000
## [19]  0.0000  0.0000  0.0000  0.0000  0.0000  0.1329  0.0650  0.0000  0.0000
## [28]  0.0509  0.1065  0.0000  0.0000
```

```
## merging atlas and data by 'atlas', 'type', 'hemi', 'side', 'roi'
```

###### d. Williams with TBV

```
##  [1] -0.08206  0.00000  0.00000  0.00000  0.00000  0.05997  0.00000 -0.11270
##  [9]  0.00000 -0.15145  0.00000 -0.07085  0.00000  0.00000  0.00000  0.00000
## [17] -0.07517  0.00000  0.00000  0.00000  0.00000  0.00000  0.00000  0.12238
## [25]  0.00000  0.00000  0.00000  0.00000  0.09578  0.00000  0.00000
```

```
## merging atlas and data by 'atlas', 'type', 'hemi', 'side', 'roi'
```

##### 2. Surface Areas

###### a. Williams no L+R and Global AI

```
## merging atlas and data by 'atlas', 'type', 'hemi', 'side', 'roi'
```

###### b. Kong no L+R and Global AI

```
## merging atlas and data by 'atlas', 'type', 'hemi', 'side', 'roi'
```

```
## Warning: Some data not merged properly. Check for naming errors in data:
##   atlas type  hemi  side  region label roi   Sex_Estimate1 Region     pval Study
##   <chr> <chr> <chr> <chr> <chr>  <chr> <chr>         <dbl> <chr>     <dbl> <chr>
## 1 <NA>  <NA>  <NA>  <NA>  <NA>   <NA>  <NA>       -0.00980 bankss… 5.65e-1 DK_K…
## 2 <NA>  <NA>  <NA>  <NA>  <NA>   <NA>  <NA>       -0.0406  fronta… 1.31e-2 DK_K…
## 3 <NA>  <NA>  <NA>  <NA>  <NA>   <NA>  <NA>        0.0638  tempor… 5.15e-4 DK_K…
## # … with 5 more variables: Sex_Estimate <dbl>, region.x <chr>, AI <dbl>,
## #   region.y <chr>, geometry <MULTIPOLYGON>
```

###### c. Williams with L+R and Global AI

```
##  [1]  0.0000  0.0000  0.0000  0.0000  0.1509  0.2356  0.0000 -0.1023  0.0830
## [10]  0.0969 -0.0960  0.0000  0.0000  0.1063  0.0802  0.0000  0.0000  0.0000
## [19]  0.0000  0.0000  0.0000  0.0000  0.0000  0.0000  0.0000  0.0000  0.0766
## [28] -0.2607 -0.1956 -0.1200  0.0820
```

```
## merging atlas and data by 'atlas', 'type', 'hemi', 'side', 'roi'
```

###### d. Williams with TBV

```
##  [1]  0.0000  0.0000  0.0000  0.0674  0.1824  0.2596  0.0787 -0.0906  0.1360
## [10]  0.1322  0.0000  0.0000  0.0000  0.1374  0.0722  0.0000  0.0000  0.0000
## [19]  0.0000  0.0000  0.0000  0.0000  0.0000  0.0000  0.0000  0.0000  0.0920
## [28] -0.2582 -0.2065  0.0000  0.1058
```

```
## merging atlas and data by 'atlas', 'type', 'hemi', 'side', 'roi'
```

#### B) Age

##### 1. Mean Thickness

###### a. Williams no L+R and Global AI

```
##  [1]  0.03047 -0.02230  0.00000 -0.02972  0.00000  0.00000 -0.03993  0.00000
##  [9]  0.00000  0.02439  0.00000 -0.02587 -0.02966 -0.03230  0.00000  0.00000
## [17]  0.02359  0.00000  0.00000 -0.03872  0.00000 -0.02644  0.00000  0.00000
## [25]  0.00000  0.00000  0.00000 -0.03009 -0.02240 -0.02776  0.00000
```

```
##  [1] -0.0291  0.0000 -0.0642  0.0000  0.0000  0.0331  0.0000  0.0000  0.0000
## [10] -0.0627 -0.0572 -0.0328  0.0000  0.0000  0.0000  0.0000  0.0000  0.0000
## [19] -0.0728  0.0000  0.0000  0.0000 -0.0237 -0.0529 -0.0269 -0.0376  0.0000
## [28]  0.0274  0.0000  0.0000  0.0000
```

```
##  [1] -0.01640062  0.00000000  0.00000000  0.00000000  0.02904376  0.00000000
##  [7]  0.00000000  0.00000000  0.00000000  0.00000000  0.00000000  0.00000000
## [13]  0.00000000  0.00000000  0.00000000  0.00000000  0.00000000  0.00000000
## [19]  0.00000000  0.00000000  0.00000000  0.00000000  0.00000000  0.00000000
## [25]  0.00000000  0.00000000  0.00000000  0.00000000  0.00000000  0.00000000
## [31]  0.00000000  0.00000000  0.00000000  0.00000000
```

```
##  [1] 0.0000000 0.0000000 0.0000000 0.0000000 0.0000000 0.0000000 0.0000000
##  [8] 0.0000000 0.0000000 0.0000000 0.0000000 0.0000000 0.0000000 0.0000000
## [15] 0.0000000 0.0000000 0.0000000 0.0000000 0.0000000 0.0000000 0.0000000
## [22] 0.0000000 0.0000000 0.0000000 0.0000000 0.0000000 0.0000000 0.0000000
## [29] 0.0225976 0.0000000 0.0000000 0.0000000 0.0000000 0.0000000
```

```
## merging atlas and data by 'atlas', 'type', 'hemi', 'side', 'roi'
```

###### b. Kong no L+R and Global AI

```
## merging atlas and data by 'atlas', 'type', 'hemi', 'side', 'roi'
```

```
## Warning: Some data not merged properly. Check for naming errors in data:
##   atlas type  hemi  side  region label roi   Age_Estimate1 Region     pval Study
##   <chr> <chr> <chr> <chr> <chr>  <chr> <chr>         <dbl> <chr>     <dbl> <chr>
## 1 <NA>  <NA>  <NA>  <NA>  <NA>   <NA>  <NA>        0.00140 bankssts  0.866 DK_K…
## 2 <NA>  <NA>  <NA>  <NA>  <NA>   <NA>  <NA>        0.00774 frontalp… 0.410 DK_K…
## 3 <NA>  <NA>  <NA>  <NA>  <NA>   <NA>  <NA>       -0.00993 temporal… 0.221 DK_K…
## # … with 5 more variables: Age_Estimate <dbl>, region.x <chr>, AI <dbl>,
## #   region.y <chr>, geometry <MULTIPOLYGON>
```

###### c. Williams with L+R and Global AI

```
##  [1]  0.0000  0.0000 -0.0551  0.0000  0.0000  0.0270  0.0000  0.0000  0.0000
## [10] -0.0577 -0.0528 -0.0268  0.0000  0.0000  0.0000  0.0000  0.0000  0.0000
## [19] -0.0633  0.0000  0.0000  0.0000  0.0000 -0.0385  0.0000 -0.0160  0.0000
## [28]  0.0361  0.0000  0.0293  0.0000
```

```
## merging atlas and data by 'atlas', 'type', 'hemi', 'side', 'roi'
```

###### d. Williams with TBV

```
##  [1] -0.0291  0.0000 -0.0642  0.0000  0.0000  0.0331  0.0000  0.0000  0.0000
## [10] -0.0627 -0.0572 -0.0328  0.0000  0.0000  0.0000  0.0000  0.0000  0.0000
## [19] -0.0728  0.0000  0.0000  0.0000 -0.0237 -0.0529 -0.0269 -0.0376  0.0000
## [28]  0.0274  0.0000  0.0000  0.0000
```

```
## merging atlas and data by 'atlas', 'type', 'hemi', 'side', 'roi'
```

##### 2. Surface Areas

###### a. Williams no L+R and Global AI

```
## merging atlas and data by 'atlas', 'type', 'hemi', 'side', 'roi'
```

###### b. Kong no L+R and Global AI

```
## merging atlas and data by 'atlas', 'type', 'hemi', 'side', 'roi'
```

```
## Warning: Some data not merged properly. Check for naming errors in data:
##   atlas type  hemi  side  region label roi   Age_Estimate1 Region     pval Study
##   <chr> <chr> <chr> <chr> <chr>  <chr> <chr>         <dbl> <chr>     <dbl> <chr>
## 1 <NA>  <NA>  <NA>  <NA>  <NA>   <NA>  <NA>       -0.0164  bankssts 0.0492 DK_K…
## 2 <NA>  <NA>  <NA>  <NA>  <NA>   <NA>  <NA>       -0.00763 frontal… 0.401  DK_K…
## 3 <NA>  <NA>  <NA>  <NA>  <NA>   <NA>  <NA>        0.0120  tempora… 0.196  DK_K…
## # … with 5 more variables: Age_Estimate <dbl>, region.x <chr>, AI <dbl>,
## #   region.y <chr>, geometry <MULTIPOLYGON>
```

###### c. Williams with L+R and Global AI

```
##  [1]  0.03047 -0.02230  0.00000 -0.02972  0.00000  0.00000 -0.03993  0.00000
##  [9]  0.00000  0.02439  0.00000 -0.02587 -0.02966 -0.03230  0.00000  0.00000
## [17]  0.02359  0.00000  0.00000 -0.03872  0.00000 -0.02644  0.00000  0.00000
## [25]  0.00000  0.00000  0.00000 -0.03009 -0.02240 -0.02776  0.00000
```

```
## merging atlas and data by 'atlas', 'type', 'hemi', 'side', 'roi'
```

###### d. Williams with TBV

```
##  [1]  0.035 -0.027  0.000 -0.025  0.000  0.000 -0.038  0.000  0.000  0.000
## [11] -0.024 -0.026  0.000 -0.025  0.000  0.000  0.000  0.000  0.000 -0.040
## [21]  0.000 -0.028  0.000  0.000  0.000  0.000  0.000 -0.027  0.000 -0.029
## [31]  0.027
```
